# Olfactory-driven beta band entrainment of limbic circuitry during neonatal development

**DOI:** 10.1101/2021.10.04.463041

**Authors:** Johanna K. Kostka, Ileana L. Hanganu-Opatz

## Abstract

Cognitive processing relies on the functional refinement of the limbic circuitry during the first two weeks of life. During this developmental period, when most sensory systems are still immature, the sense of olfaction acts as “door to the world”, providing the main source of environmental inputs. However, it is unknown whether early olfactory processing shapes the activity in the limbic circuitry during neonatal development. Here, we address this question by combining simultaneous *in vivo* recordings from the olfactory bulb (OB), lateral entorhinal cortex (LEC), hippocampus (HP), and prefrontal cortex (PFC) with olfactory stimulation as well as opto- and chemogenetic manipulations of mitral/tufted cells (M/TCs) in the OB of non-anesthetized neonatal mice. We show that the neonatal OB synchronizes the limbic circuity in the beta frequency range. Moreover, it drives neuronal and network activity in LEC, as well as subsequently, HP and PFC via long-range projections from mitral cells (MCs) to HP-projecting LEC neurons. Thus, OB activity shapes the communications within limbic circuits during neonatal development.

## INTRODUCTION

Coordinated neuronal activity during early development refines the neural circuits that account for complex processing in the adult brain. During the first two postnatal weeks, when the eyes and ears of rodents are still closed and they perform no active whisking, coordinated activity patterns in the sensory periphery occur mainly independently of sensory input [1,2]. Even if strong auditory and visual stimuli can evoke activity in the corresponding primary sensory cortices before eye and ear opening [3-6] and sensory feedback evoked by twitches during active sleep promotes cascades of neuronal activity in the primary somatosensory cortex [7], prominent activity patterns in the developing primary sensory cortices are largely driven by spontaneous neuronal discharges in the immature peripheral sensory structures [8]. Both spontaneous and stimulus-driven activity trigger discontinuous oscillatory bursts in the corresponding primary sensory cortices [5,9–12] that are necessasry for the development of sensory discrimination [13]. Similar activity patterns can also be observed in cortical areas involved in higher cognitive processing. During active sleep, sensory feedback-evoked activity in the primary somatosensory cortex entrains the HP via the medial entorhinal cortex in beta oscillations and excitatory activity of entorhinal cortex neurons instructs hippocampal development during postnatal ages [14,15,16]. Moreover, neonatal hippocampal sharp-waves are often triggered by twitches but also occur when input from the brainstem is lesioned [17]. Further, discontinuous theta band oscillations in the LEC drive similar activity patterns in the HP, which in turn entrains the prelimbic area (PL) of the PFC [18-21]. Disturbance of these early activity patterns in mouse models of psychiatric risk [22-26] as well as through pharmacological [27] or optogenetic manipulations [28] led to disruption of adult circuits and behavioral abilities.

Due to the limited or absent functionality of the visual and auditory systems during the first two postnatal weeks, their contribution to the development of limbic networks has been considered negligible. This hypothesis has been supported by data showing that the synchrony between V1 and the HP-PFC network before eye-opening is rather weak [20]. Besides reafferent signals evoked by twitches, neonatal mice process olfactory inputs from birth on and use this information to instruct learning and cue-directed behaviors essential for their survival [29]. Correspondingly, the anatomical pathways from OB to cortical areas are unique among sensory systems as they lack the relay via the thalamus. MCs send afferents directly to the piriform cortex (PiR) and limbic brain areas such as LEC and amygdala [30,31]. At adult age, in line with the anatomical connectivity, strong functional coupling during odor processing has been found between OB and these brain areas. For example, adult olfactory processing relies on respiration-modulated beta and gamma OB activity [32-34]. Further, beta oscillations in PiR, LEC and HP play a critical role in olfactory memory processing [35-37]. Moreover, synchronized beta oscillations between OB-HP and LEC-HP are critically involved in odor learning [38-41]. Recently, beta oscillations in prefrontal-hippocampal networks have been identified to support the utilization of odor cues for memory-guided decision-making [42].

The tight and behaviorally relevant coupling between OB and limbic circuits at adult age leads to the question, which role does olfactory activation early in life play for these circuits. Previously, we showed that discontinuous oscillatory activity in theta-beta range, emerging as a result of bursting MCs in the neonatal OB, entrains similar oscillatory patterns in the LEC [43,44] of anesthetized mice. However, it remains unknown whether a similar entrainment occurs in a non-anesthetized state. Furthermore, the role of neuronal and network activity in the OB for the functional entrainment of downstream areas within the neonatal hippocampal-prefrontal network is still largely unknown.

To address these knowledge gaps, we simultaneously monitored single-unit activity (SUA) and local field potentials (LFP) in OB, LEC, HP, and PFC of non-anesthetized neonatal mice (postnatal day (P) 8-10) during odor stimulation and manipulation of M/TC activity using excitatory opsins and inhibitory DREADDs. We show that odor stimulation and optogenetic activation of M/TCs triggers action potential firing in LEC and HP as well as prominent beta oscillations that synchronize the OB with the downstream cortical areas. Conversely, blocking MC output specifically diminishes OB - limbic network coupling in the beta frequency range.

These data document the ability of coordinated activity at the sensory periphery of newborns to shape the network activity in circuits accounting for adult cognitive processing.

## RESULTS

### Oscillatory activity in OB times the network activity in limbic circuits of neonatal mice

To get first insights into the impact of OB activity on developing cortical circuits including LEC, HP, and PFC, we simultaneously recorded the LFP and SUA in all four brain areas in non-anesthetized neonatal (P8-10) mice (n=56, Figs 1A and 1B) and assessed the temporal relationships between network oscillations and neuronal firing. All investigated areas showed discontinuous oscillatory activity in the theta-beta range [20,21,43], accompanied by continuous low amplitude slow frequency oscillations peaking at 2-4 Hz (respiration rhythm, RR). Half of the oscillatory events detected in OB (median (med): 53.8 %, interquartile range (iqr): 47.7 – 65.1 %, n=20) co-occurred in all four brain regions. To quantify the coupling of OB to cortical areas, we calculated the imaginary coherence (Fig 1C). While a high level of synchrony linked OB with all investigated cortical areas, the strength of coupling was frequency-dependent, having the highest magnitude in the beta frequency range for OB-LEC and OB-HP and in the RR frequency band for OB-LEC and OB-PFC (Fig 1C). The beta band imaginary coherence between OB and LEC was higher when compared with the previously reported values in anesthetized mice [43]. Anesthesia shifts the OB-LEC synchronization towards lower frequencies.

**Fig 1.**
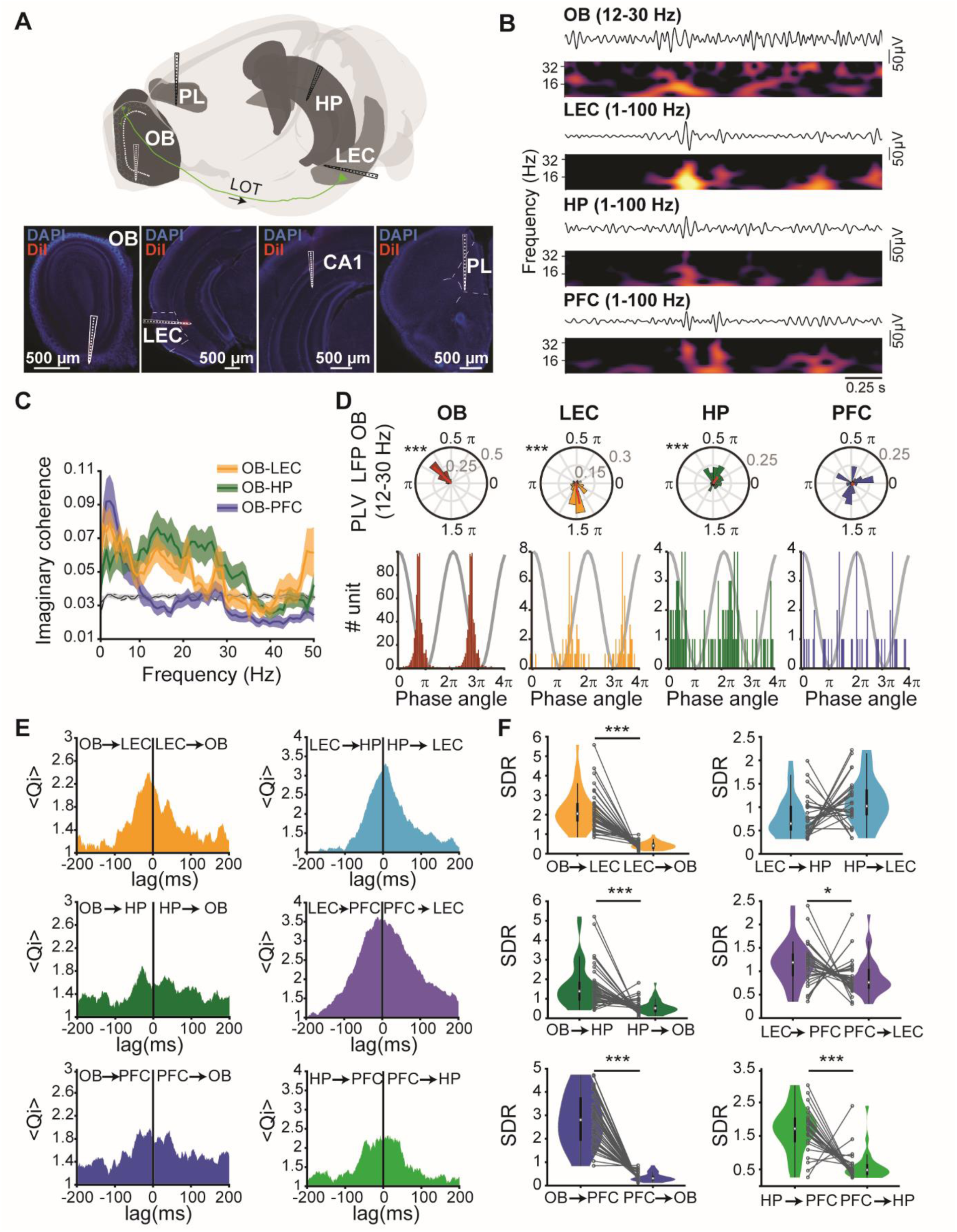
Functional coupling between neonatal OB, LEC, HP, and PFC. **(A)** Top, schematic of recording configuration for simultaneous extracellular recordings in OB, LEC, HP, and PFC. The positions of recording sites were displayed superimposed on the corresponding brain areas (Brainrender: [48]). Bottom, digital photomontages displaying the DiI-labeled (red) electrode tracks in DAPI (blue) stained slices of OB, LEC, HP, and PL of a P10 mouse. **(B)** Representative beta band filtered LFP traces recorded in the OB, LEC, HP, and PFC displayed together with the wavelet spectra of beta oscillations recorded simultaneously. **(C)** Mean spectra of imaginary coherence calculated for OB-LEC (yellow, n= 42), OB-HP (green, n= 38), and OB-PFC (blue, n= 40). The gray lines correspond to the significance threshold as assessed by Monte Carlo simulation. Bounded lines correspond to the SEM. **(D)** Top, polar plots displaying the phase-locking of significantly locked units in OB (red), LEC (yellow), HP (green), and PFC (blue) to beta oscillations in OB. Bottom, histograms of mean phase angle for significantly phase-locked OB (red), LEC (yellow), HP (green), and PFC (blue) units. Histograms are replicated over two OB beta cycles (gray curve). (*** p < 0.001, Rayleigh test for non-uniformity). **(E)** Plots of standardized mean spike-spike cross-covariance for OB-LEC (yellow), OB-HP (green), OB-PFC (blue), LEC-HP (light blue), LEC-PFC (purple), and HP-LEC (light green). Negative lags indicate that spiking in the first brain area precedes spiking in the second brain area. **(F)** Spectral dependency ratio (SDR) calculated for OB-LEC (yellow), OB-HP (green), OB-PFC (blue), LEC-HP (light blue), LEC-PFC (purple), and HP-LEC (light green). Gray dots and lines correspond to individual animals. (* p < 0.05, ** p < 0.01, *** p < 0.001, Wilcoxon signed-rank test).

To uncover whether OB activity times the neuronal firing of cortical areas, we calculated the phase-locking of SUA recorded in OB, LEC, HP, and PFC to beta band oscillations (12-30 Hz) in OB (Fig 1D). Significantly locked OB units fired shortly before the trough of the beta cycle, while LEC and HP units were locked to significantly shifted phase angles (Figs 1D, S1, and S2 Tables). In contrast, units in LEC and HP fire preferentially at the trough of local entorhinal and hippocampal beta oscillations, respectively. This locking pattern was detected for both significantly locked and not significantly locked units to the OB beta phase (S1A Fig).

These data suggest a phase shift from OB to LEC and HP. Solely the prefrontal firing showed no phase preference of locking to the oscillatory phase in OB. Next, we questioned whether the communication between OB and cortical areas is directed and whether OB acts as a driving force within the circuit. For this, we assessed the temporal relationship between the firing in cortical regions and OB by calculating the standardized cross-covariance of unit pairs [45]. For unit pairs between OB and LEC, OB and HP the peak of cross-covariance was at negative time-lags, indicating that spiking in OB preceded cortical firing (Fig 1E). Monitoring the timing of interactions between cortical areas (LEC-HP, LEC-PFC, and HP-PFC) confirmed the previously reported directionality of communication [21], yet less clear as for the OB-driven coupling. The cross-covariance of unit pairs OB-LEC, OB-HP, LEC-HP, LEC-PFC, and HP-PFC predominantly peak at negative lags (OB-LEC: negative: 60.3%, positive: 39.7%; OB-HP: negative: 51.2%, positive: 48.8%, LEC-HP: negative: 50.8 %, positive: 49%; LEC-PFC: 52.5%, positive: 47.5%; HP-PFC: negative: 56.2%, positive: 43.8%). Solely more unit pairs between OB and PFC showed more peak positive lags compared to peak negative lags (negative: 48.2%, positive: 51.8%) (S1B Fig). As spike-dependent methods are strongly biased by the firing rate of investigated neurons, which is rather low in neonatal mice, we next used the spectral dependency ratio (SDR), a method that infers causal direction from time-series data [46,47], to confirm the directed communication between OB and cortical areas. SDR values for OB → LEC were significantly higher than for LEC → OB, supporting the drive from OB to LEC. Further, the SDR analysis revealed a spectral dependency of HP as well as PFC on OB, suggesting the contribution of OB activity to the oscillatory entrainment of prefrontal and hippocampal circuits (Fig 1F and S3 Table). Moreover, the analysis confirmed the previously reported directed interaction from HP to PFC and LEC to PFC [20,21]. No SDR difference was detected for LEC-HP, indicating that, in line with anatomical data [21], a bidirectional coupling links HP and LEC (Fig 1F and S3 Table).

Thus, tight directed interactions between OB and cortical areas ensure timed firing and oscillatory entrainment within downstream LEC-HP circuits.

### Olfactory stimulation drives firing and network synchronization along the OB-LEC-HP-PFC pathway

In adult mice, olfactory sampling has been reported to induce firing and augment the synchrony within LEC-HP-PFC networks [34–38,42]. Even in anesthetized neonatal mice, odor exposure led to an increase in beta band activity in LEC [43]. However, it is unknown, which impact anesthesia has on these effects and whether odor sampling engages also the downstream areas, HP and PFC, in neonatal mice. To address these questions, LFP and SUA were recorded in the OB, LEC, HP, and PFC of neonatal mice (n=19) before, during and after stimulation (2 s) with up to four different neutral odors (Fig 2A). Odor stimulation activated 49,9% of OB units (Fig 2B and 2C). While most OB units (49,9%) responded with an increase in their firing rate, 13,2% of them were inhibited in response to the odor stimulation (Fig 2C). The augmented firing has been detected also in downstream areas, yet at a different latency from the first inhalation during odor exposure. Units in LEC responded faster than units in HP and LEC, the activation of which was long-lasting and spanned several inhalation cycles (Fig 2B and 2D). The percentage of significantly activated units was higher for LEC (24.5%) and PFC (18.8%) when compared to HP (6.8%), where also significantly inhibited units (7.3%) in response to odor stimulation have been detected (Fig 2C). Odor stimulation broadly (2-40 Hz) increased the oscillatory power in all four brain areas (Fig 2E and S4 Table) and the synchrony in RR and beta band between OB and LEC, OB and HP as well as OB and PFC (Fig 2F and S5 Table). The observed increase in power and synchrony was independent of the specific odor that was presented. These data demonstrate that odor exposure drives the oscillatory coupling of OB and downstream limbic regions already at neonatal age.

**Fig 2.**
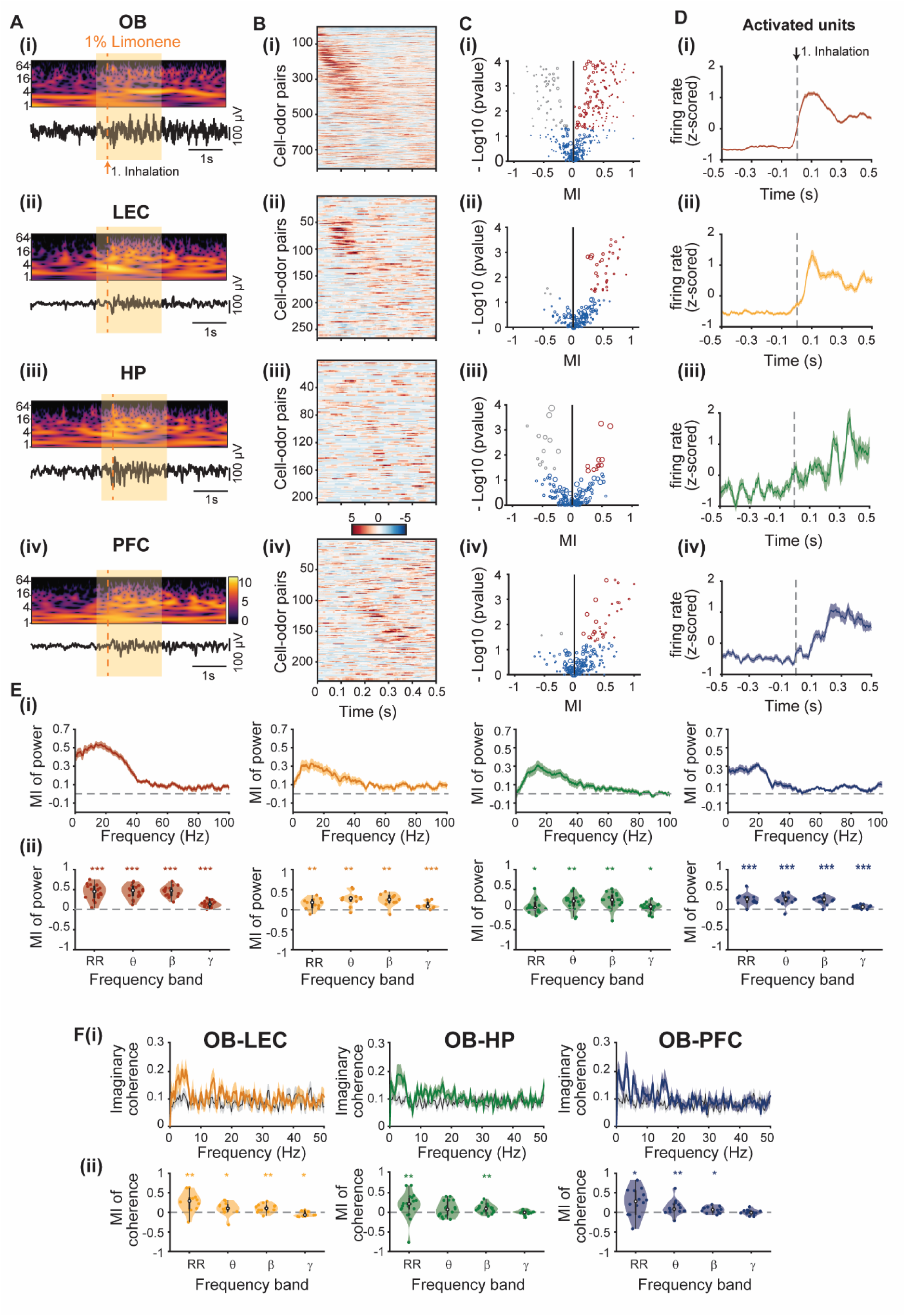
Effects of olfactory stimulation on firing and oscillatory activity in OB, LEC, HP, and PFC. **(A)** Representative LFP trace extracellularly recorded in the **(i)** OB, **(ii)** LEC, **(iii)** HP, and **(iv)** PFC during stimulation with 1% limonene accompanied by the corresponding wavelet spectrum. **(B)** Plots of odor-evoked firing rates for cell-odor pairs in **(i)** OB, **(ii)** LEC, **(iii)** HP, and **(iv)** PFC aligned to the first inhalation during odor stimulation and ordered by peak firing time. **(C)** MI of SUA of cell-odor pairs in response to odor stimulation of **(i)** OB, **(ii)** LEC, **(iii)** HP, and **(iv)** PFC. Significantly activated units are shown in red, whereas significantly inhibited units in gray, p < 0.05, Wilcoxon signed-rank test. **(D)** Z-scored firing rate of activated cell-odor pairs in **(i)** OB (red), **(ii)** LEC (yellow), **(iii)** HP (green), and **(iv)** PFC (blue) in response to the first inhalation during odor stimulation. **(E) (i)** Plot of MI for power during stimulation with 1% limonene for OB (red), LEC (yellow), HP (green), and PFC (blue). **(ii)** Mean MI of LFP power in different frequency bands for OB (red), LEC (yellow), HP (green), and PFC (blue). (* p < 0.05, ** p < 0.01, *** p < 0.001, Wilcoxon signed-rank test). **(F) (i)** Plot of MI for coherence during stimulation with 1% limonene for OB-LEC (yellow), OB-HP (green), and OB-PFC (blue). **(ii)** Mean MI of coherence in different frequency bands for OB-LEC (yellow), OB-HP (green), and OB-PFC (blue) (* p < 0.05, ** p < 0.01, *** p < 0.001, Wilcoxon signed-rank test).

### Activation of M/TCs induces beta oscillations in neonatal OB

To elucidate the mechanisms of directed communication between OB and downstream cortical areas, we activated ChR2-transfected M/TCs by light and simultaneously monitored the network and neuronal activity in neonatal LEC, HP, and PFC. Transfection of M/TCs was achieved using a cre-dependent virus vector (AAV9-Ef1a-DIO-hChR2(E123T/T159C)-EYFP) that was injected into the right OB of P1 Tbet-cre mice (Fig 3A). ChR2-EYFP expression was reliably detected in M/TCs and their projections 7 days after injection (Fig 3B). Ramp light stimuli of increasing intensity (473 nm, total duration 3 s) were used to activate M/TCs in the OB of P8-10 mice (Fig 3A). The stimulation parameters have been set in line with previous data [49] to prevent not only firing as a result of tissue heating but also artificially synchronous firing patterns and large stimulation artifacts. Ramp stimulation led to a sustained increase of spike discharge and broad-band (4-100 Hz) LFP power augmentation in OB that peaked in the beta frequency range (12-30 Hz) (Figs 3C, 3D, and S2A Fig). Light-evoked beta oscillations showed current-source density (CSD) profiles with alternating sinks and sources over MCL and low EPL, respectively (S2C Fig). They are similar to the CSD profile of OB beta oscillations induced by passive sniffing in neonatal (S2C Fig) and adult mice [50]. In cre^+^ mice, the modulation indices (MI) for theta, beta, and gamma power were significantly increased and different from those calculated for cre^-^ animals (Fig 3Dii and S6 Table). Correspondingly, SUA is strongly augmented during ramp stimulation (Figs 3E and 3F). This activation was not layer-specific and, mirroring the tight OB wiring, not only M/TCs but also granule cells (GCs) and other OB interneurons increased their firing in response to light activation of ChR2-transfected M/TCs (S2B and S2D Figs). Units recorded in the GCL had shorter halfwidths and smaller amplitudes when compared to units recorded in MCL, (S2E and S2F Figs). The different waveform suggests that these units are most likely GCs and not MCs. Analysis of the firing onset along OB layers confirmed the global activation. Cells in the MCL and GCL started to fire immediately after the 3 ms-long light pulses, whereas cells in the extra plexiform layer (EPL) and glomerular layer (GL) responded with a brief delay (Fig 3G).

**Fig 3.**
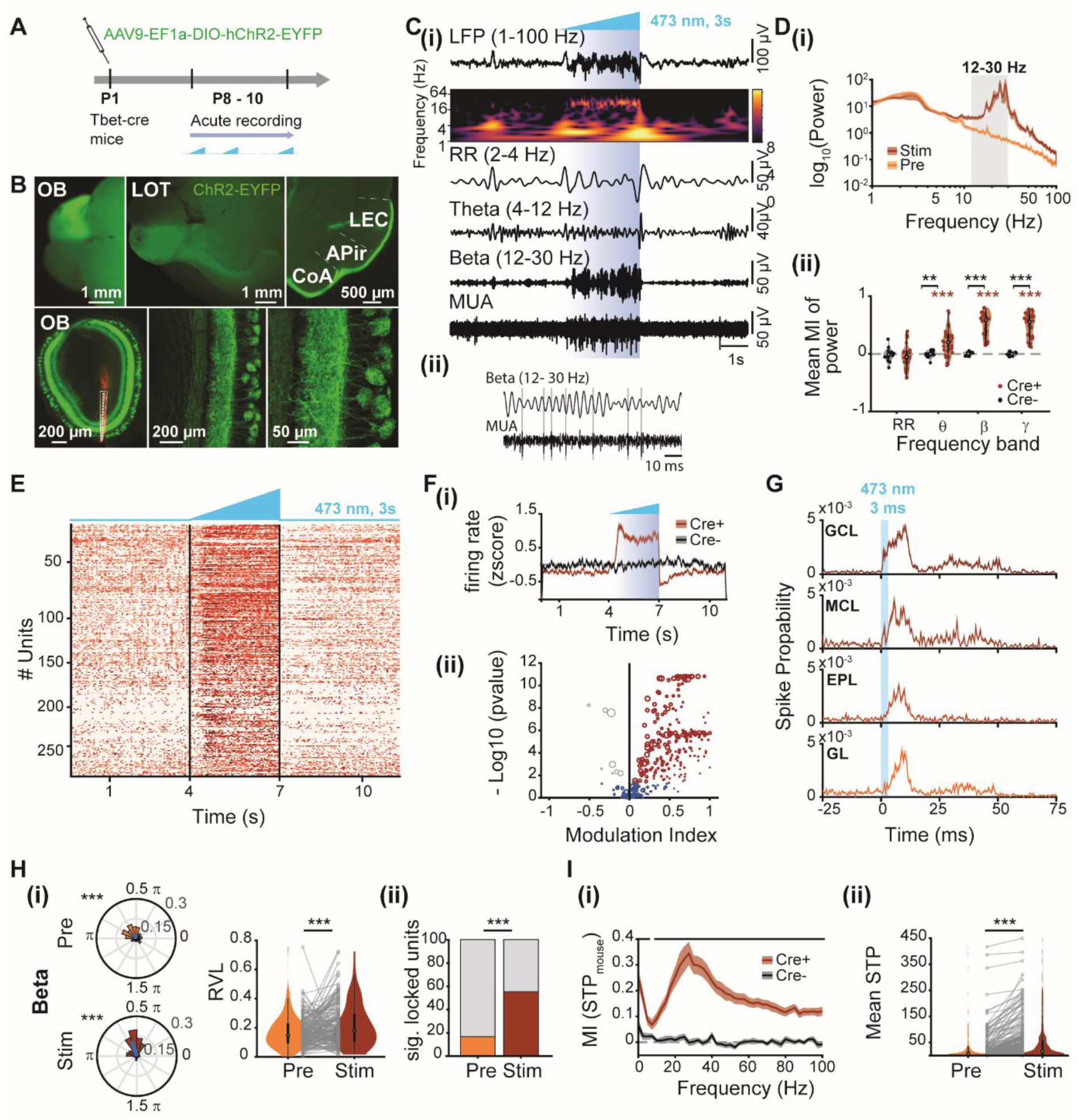
Effects of M/TC manipulation by light on single-unit entrainment and oscillatory activity in OB. **(A)** Schematic of the experimental protocol. **(B)** Top, photograph of the dorsal (left) and ventral side (middle) of a brain from a Tbet-cre^+^ mouse showing EYFP expression in the OB and M/TC axonal projections (LOT) to LEC, piriform transition area (APir), and cortical amygdala (CoA) (right). Bottom, digital photomontages displaying the DiI labeled electrode track in OB (left) and confocal images displaying the mitral cell layer (MCL) of the right OB at different magnifications (middle and right). **(C)**(i) Representative extracellularly recorded LFP in the OB displayed band-pass filtered in different frequency bands and accompanied by the corresponding wavelet spectrum during ramp stimulation, as well as by the simultaneously recorded MUA in the MCL. (ii) Magnification of a beta oscillation during ramp stimulation and corresponding MUA trace. Gray lines indicate times of action potential occurrence. **(D)** (i) Power spectrum for OB LFP before (orange) and during (red) ramp stimulation. The gray shaded area corresponds to the beta band (12-30 Hz). (ii) Mean MI of LFP power in different frequency bands for cre^+^ (red) and cre^-^ (black) mice. (red stars for cre^+^: *** p < 0.001, Wilcoxon signed-rank test; black stars for comparison cre^+^ vs. cre^-^: ** p < 0.01, *** p < 0.001, Wilcoxon rank-sum test). **(E)** Raster plot of SUA in the OB before, during, and after ramp stimulation. **(F) (i)** Z-scored firing rate in response to ramp stimulation of units recorded in the OB of cre^+^ (red) and cre^-^ (black) mice. **(ii)** MI of SUA firing in response to ramp stimulation (Significantly activated units are shown in red, whereas significantly inhibited units in gray, p < 0.01, Wilcoxon signed-rank test). **(G)** Spiking probability of units located in the granule cell layer (GCL), MCL, external plexiform layer (EPL), and glomerular layer (GL) after a 3 ms light pulse (blue box, 473 nm) delivered to the OB. **(H) (i)** Phase locking of OB units to beta oscillations in OB. Left, polar plots displaying phase locking of OB units before (Pre, orange) and during ramp stimulation (Stim, red). The mean resulting vectors are shown as blue lines. (*** p < 0.001, Rayleigh test for non-uniformity). Right, violin plots displaying the resulting vector length (RVL) of OB units before (Pre, orange) and during ramp stimulation (Stim, red). Gray dots and lines correspond to individual units. (*** p < 0.001, linear mixed-effect model). **(ii)** Percentage of significantly locked units before (Pre, yellow) and during (Stim, red) stimulation. (*** p < 0.001, Fisher’s exact test). **(I) (i)** Plot of mean MI of spike-triggered power (STP) for cre^+^ (red) and cre^-^ (black) mice during ramp stimulation. (black line: p < 0.05, Wilcoxon rank-sum test). **(ii)** Violin plots displaying mean STP for OB units before (Pre, yellow) and during ramp stimulation (Stim, red). Gray dots and lines correspond to individual units. (*** p < 0.001, linear mixed-effect model).

To assess the temporal relationship between neuronal firing and beta oscillations in OB, we calculated the locking of SUA firing to the oscillatory phase before (Pre) and during (Stim) light stimulation (Fig 3Cii). Ramp stimulation caused a significantly stronger locking of OB units to beta oscillations (Pre: med: 0.147, iqr: 0.094 – 0.227; Stim: med: 0.180, iqr: 0.103 – 0.294, n_units_=176 from 26 mice, p=9.39*10^-5^, LMEM) (Fig 3Hi) and an augmentation of the proportion of significantly phase-locked units to the beta rhythm during ramp stimulation (Pre: 16.5 %, 29/176 units, Stim: 55.1 %, 97/176 units, p=3.12*10^-14^, Fisher’s exact test) (Fig 3Hii). Of note, the coupling of OB units to the RR phase was weaker (Pre: med: 0.129, iqr: 0.080 – 0.220, Stim: med: 0.097, iqr: 0.057 – 0.157, n_units_=176 from 26 mice, p=7.782*10^-6^, LMEM) (S3Ai and S3Aii Figs) even though the proportion of locked units (Pre: 14.8 %, 26/176 units, Stim: 14.2 %, 25/176 units, p=1, Fisher’s exact test) and the power of RR oscillations were not altered upon light stimulation (Fig 3Dii and S3Aii Figs). In contrast, light stimulation had no effects on the phase-locking of OB units to oscillatory phase in cre^-^ mice (S3B Fig and S7 Table). The larger beta power observed during ramp stimulation might result from increased M/TC and interneuronal firing, since spike-triggered power (STP) analysis revealed that the ability of OB units to trigger beta power is stronger during ramp stimulation compared to baseline periods (Pre: med: 6.694 µV², iqr: 2.291 – 16.447 µV²; Stim: med: 18.285 µV², iqr: 4.437 – 58.407 µV²; n_units_ = 309 from 19 mice, p = 3.16*10^-13^, LMEM) (Fig 3I). These data indicate that the activation of M/TCs recruits the local circuitry in the OB and thereby, organizes the OB network activity in the beta rhythm.

### M/TC activation drives neuronal firing in LEC and HP

To characterize the downstream effects of beta band entrainment of OB, we firstly analyzed the organization of OB projections in neonatal mice. In line with morphological investigations in adult mice [30], we previously showed that MC axons are present in superficial layers of LEC already at neonatal age [43]. Entorhinal neurons in layer II/III strongly project to HP and weakly to PFC [21,26]. Here, we performed axonal tracing of M/TCs using the anterograde virus (AAV9-hSyn-hChR2(H134R)-EYFP) injected into the OB at P8. Simultaneously, we monitored the entorhinal neurons that project to HP by using the retrograde virus (AAVrg-CamKIIa-mCherry) injected into the HP at P8 (Figs 4A and 4B). At P18, MC axons expressing EYFP were present in layer I/II of LEC and PiR (Fig 4B). Additionally, mCherry-expressing HP-projecting neurons were identified in entorhinal layer II/III. These neurons send their apical dendrites to layer I of LEC, where they collocate with MC axonal projections (Fig 4B).

**Fig 4.**
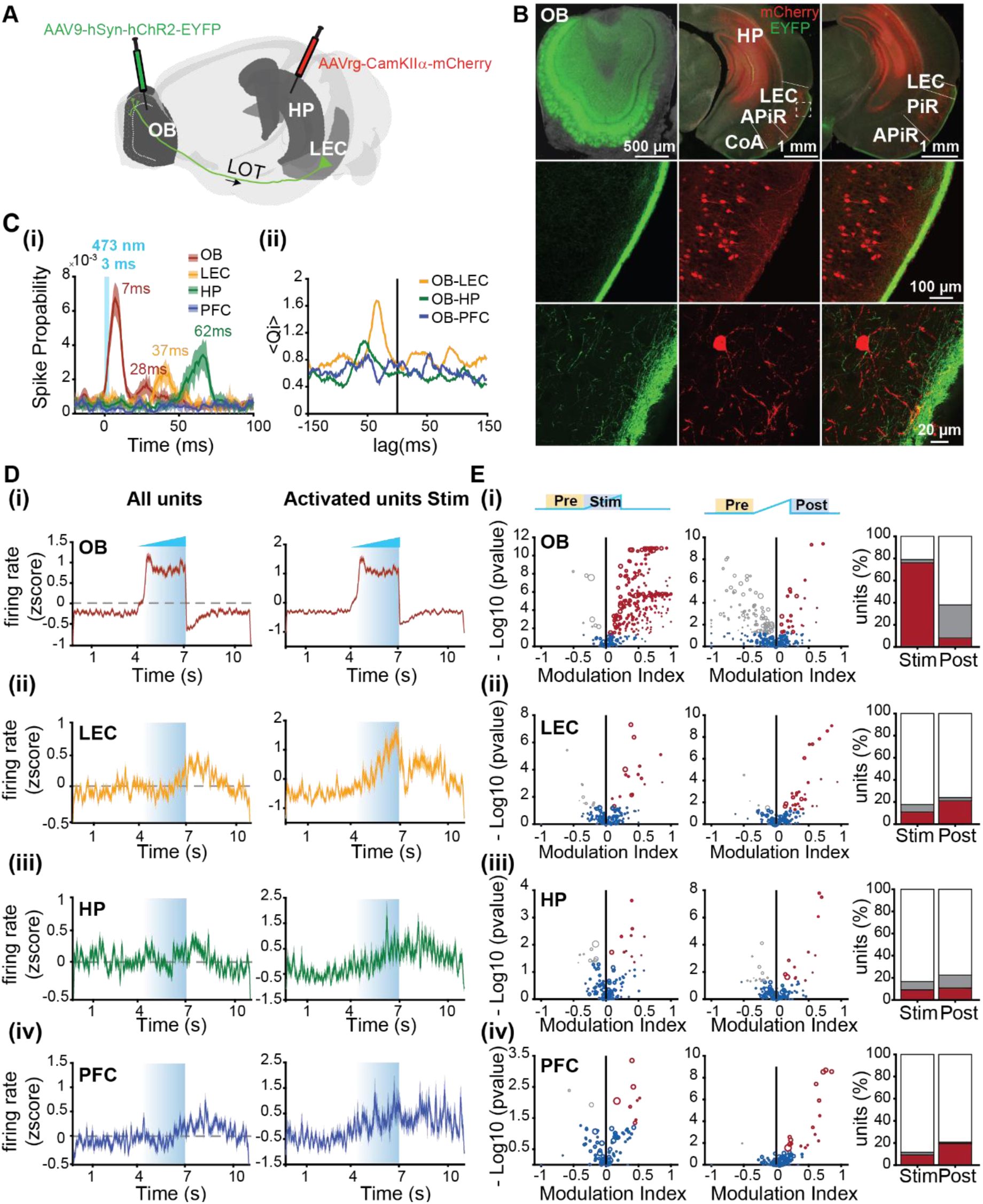
Effects of optogenetic manipulation of M/TCs on single-unit activity in LEC, HP, and PFC. **(A)** Schematic of the experimental protocol used to trace MC axons and neurons projecting to HP. (Brainrender: [48]). **(B)** Top, digital photomontages displaying EYFP (green) and mCherry (red) fluorescence in coronal slices including OB (left, injection side of AAV9-hSyn-hChR2-EYFP), HP (middle, injection site of AAVrg-CamKIIα-mCherry), and LEC (right). Note the co-expression of EYFP and mCherry in LEC. Middle, EYFP (left), mCherry (middle), and their co-expression in the LEC are shown at larger magnification (dashed box). Bottom, EYFP (left), mCherry (middle), and their co-expression shown at larger magnification for a HP-projecting entorhinal neuron with dendrites targeting layer I. **(C) (i)** Spike probability of units in OB (red), LEC (yellow), HP (green), and PFC (blue) after a 3 ms light pulse (473 nm) delivered to the OB. Numbers indicate the delay of the peak spike probability for each brain area. All OB units (including directly responding GCL and MCL units as well as those subsequently activated) were considered. **(ii)** Spike-spike cross-covariance for OB-LEC (yellow), OB-HP (green), and OB-PFC (blue). Negative lags correspond to OB activity driving spiking in other brain areas. **(D) (i)** Left, z-scored firing rate of units recorded in the OB of cre^+^ (red) mice in response to light stimulation. Right, z-scored firing rate of significantly activated units during ramp stimulation. **(ii)** Same as (i) for units recorded in LEC (yellow). **(iii)** Same as (i) for units recorded in HP (green). **(iv)** Same as (i) for units recorded in PFC (blue). **(E) (i)** Left, volcano plot displaying the MI of SUA firing rates recorded in the OB before (Pre) vs. during (Stim) ramp stimulation (significant activated units are shown in red and significant inhibited units in gray, p < 0.01, Wilcoxon signed-rank test). Middle, same as the left image but for SUA firing rates before (Pre) vs. after (Post) ramp stimulation. Right, bar plots depicting the percentage of activated (red) and inhibited (gray) units during (Stim) and after (Post) ramp stimulation. **(ii)** Same as (i) for units recorded in LEC. **(iii)** Same as (i) for units recorded in HP. **(iv)** Same as (i) for units recorded in PFC.

Since these morphological data suggest that the OB interacts with downstream cortical areas, in a second step, we monitored the functional impact of direct OB projections on limbic circuits. For this, we used pulse (3 ms) and ramp (3 s) blue light stimulations (473 nm) of transfected OB neurons and simultaneously recorded the neuronal activity in LEC, HP, and PFC. Pulse stimulation of M/TCs induced neuronal firing in all investigated brain areas, except PFC (Fig 4Ci). While the light-evoked OB firing rate sharply peaked already 7-8 ms post-stimulus, the responses in the other brain areas were substantially broader and delayed (37 ms in LEC, 45-60 ms in HP). A second firing increase was detected in OB after ∼28 ms and might reflect OB-internal processing or feedback activation from downstream areas. To expand on these results, we used normalized cross-covariance analysis to uncover the temporal correlations between light-evoked spike trains in the investigated brain regions. The most prominent interaction was detected for OB-LEC, with OB firing preceding the entorhinal discharges (Fig 4Cii). While having a similar directionality, the OB-HP cross-covariance peaked later and less precisely. The data gives first insights into the communication pathways relaying the information from M/TCs to LEC and subsequently, to HP.

Ramp stimulation of M/TCs evoked neuronal firing in LEC, HP, and PFC with similar dynamics: a fast increase in OB followed by a delayed spiking in LEC, and subsequently in HP and PFC. In OB, SUA abruptly increased with ramp onset (76.2 % of units activated significantly, 2.8 % units inhibited significantly, p < 0.05, Wilcoxon sign-rank test Pre vs. Post for each unit) and decreased post-stimulus (8.2 % of units activated significantly, 29.9 % of units inhibited significantly, p < 0.05, Wilcoxon sign-rank test Pre vs. Post for each unit) (Figs 4Di and 4Ei). In contrast, the average SUA firing rate in LEC, HP, and PFC showed a delayed increase starting around halfway through the ramp and continuing after the light stimulation (Figs 4Dii-iv). Analysis of the proportion of activated units during and after ramp stimulation revealed that neurons in downstream areas expressed higher firing rates also after the light was switched off (Fig 4Eii-iv), indicating that the activation of M/TCs boosted the cortical network activation. Correspondingly, this post-stimulus firing increase recruited more neurons than those activated during ramp stimulation (LEC: 11.0 % during stimulation vs. 21.2 % post-stimulus; HP: 9.2 % vs. 10.8 %; PFC: 9.2 % vs. 19.5 %). In HP, the post-stimulus network effect was not restricted to the activation of neurons but also related to the increase in the proportion of neurons that were inhibited after the ramp (7.5 % vs. 11.7 %). In all four brain regions, ramp stimulation significantly modulated the neuronal firing (OB: proportion of modulated units: 0.790, CI: [0.739 0.834]; LEC: 0.178, CI: [0.125 0.248], HP: 0.167, CI: [0.111 0.243], PFC: 0.115, CI: [0.064 0.199], statistical threshold: 0.05, CI: 95 % confidence intervals for a binomial model). The firing increase in LEC, HP and PFC was accompanied by stronger coupling of LEC and HP firing, yet not PFC firing, to OB beta oscillations after light activation of M/TCs (for LEC, Pre: med: 0.106, iqr: 0.052 – 0.194; Stim: med: 0.145, iqr: 0.073 – 0.238; n_units_ = 71 from 18 mice, p = 0.024, LMEM; for HP Pre: med: 0.124, iqr: 0.063 – 0.120, Stim: med: 0.162, iqr: 0.095 – 0.276; n_units_ = 56 from 12 mice, p= 0.006, LMER; for PFC Pre: med: 0.118, iqr: 0.063 – 0.214; Stim: med: 0.126, iqr: 0.078 – 0.239; n_units_ = 59 from 13 mice, p= 0.839, LMER) (S3C, S3D and S3E Figs). In all four investigated brain regions, light stimulation of control animals did not change the proportion of modulated units when considering a 5% modulation of units as chance level (OB: proportion of modulated units: 0.009, CI: [0.002 0.03]; LEC: 0.083, CI: [0.036 0.181], HP: 0.030, CI: [0.010 0.084], PFC: 0.071, CI: [0.004 0.315], statistical threshold: 0.05, CI: 95 % confidence intervals for a binomial model) (S4A and S4B Figs).

Thus, M/TC firing drives the activation of entorhinal, and subsequently, hippocampal and prefrontal circuits.

### M/TC activation boosts beta band coupling within downstream limbic circuits

The long-lasting effects of M/TC stimulation on the neuronal firing of downstream areas, LEC, HP, and PFC suggest that OB activation might act as a driving force for the generation of network oscillations in neonatal limbic circuits. To test this hypothesis, we paired ramp light stimulation of ChR2-transfected M/TCs with LFP recordings in LEC, HP, and PFC of P8-10 mice. Ramp stimulation of M/TCs increased the oscillatory power in LEC, HP, and PFC for a broad frequency range spanning from theta to gamma oscillations (Figs 5A, 5B, and S6 Table). Cre^+^ and cre^-^ had significant different MI power values for the beta band (LEC: p=0.033, HP: p=0.021; PFC: p=2.78*10^-4^, Wilcoxon rank-sum test). Moreover, we assessed the degree of synchrony between OB and cortical areas during light stimulation by calculating the imaginary part of coherence, a measure that is insensitive to false connectivity arising from volume conduction [51]. The imaginary coherence between OB and LEC, OB and HP as well as OB and PFC increased during light activation of M/TCs, the most prominent effects being detected in the beta band range (Fig 5C and S8Table). MI values for coherence were significantly higher than 0 (OB-LEC: p=0.005, OB-HP: p=0.005; OB-PFC: p=0.0004, Wilcoxon singed-rank test) and significantly different from cre^-^ mice for the beta band for OB-LEC, OB-HP, and OB-PFC (OB-LEC: p=0.036, OB-HP: p=0.017; OB-PFC: p=0.008, Wilcoxon rank-sum test) (S8 Table).

**Fig 5.**
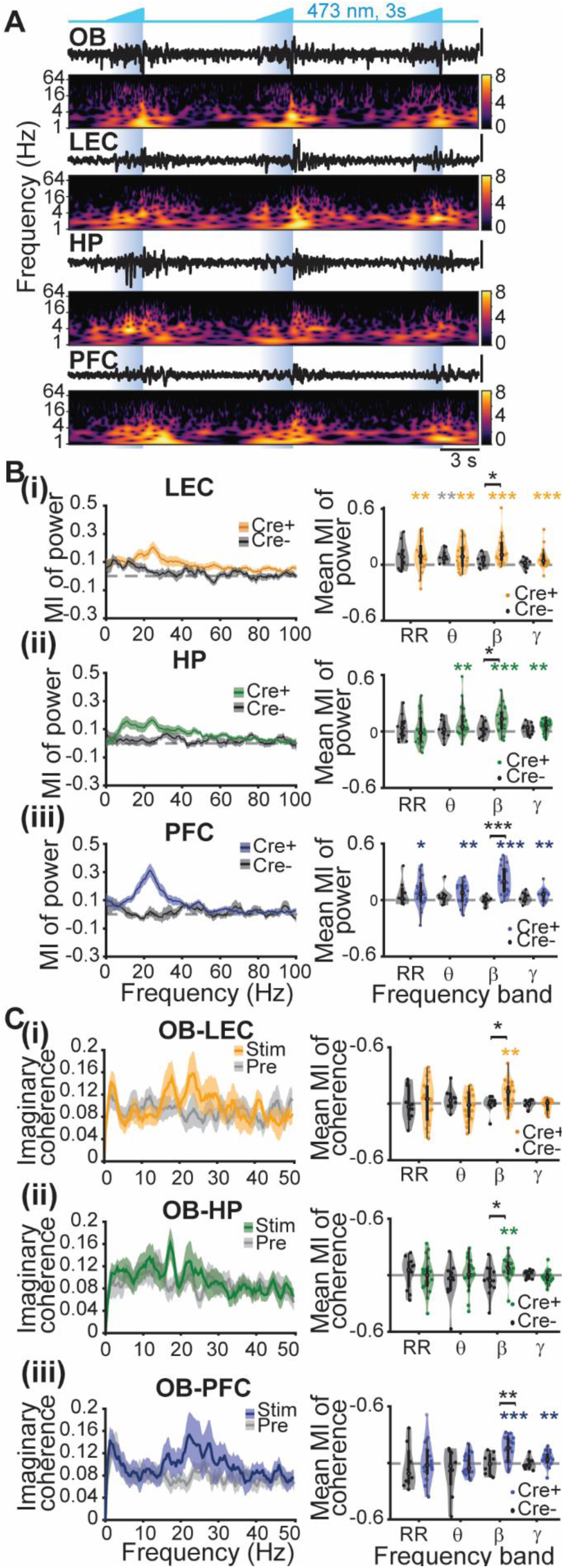
Oscillatory entrainment of limbic circuits as a result of M/TC activation by light. **(A)** Representative LFP traces recorded in the OB, LEC, HP, and PFC during ramp stimulation of ChR2-transfected M/TCs accompanied by the corresponding wavelet spectra. Vertical black lines correspond to 100 µV. **(B) (i)** Left, plot of MI for power during ramp stimulation of oscillations in LEC for cre^+^ (yellow) and cre^-^ (black) mice. Right, MI of LFP power averaged for different frequency bands for cre^+^ (yellow) and cre^-^ (black) mice. **(ii)** Same as (i) for HP. **(iii)** Same as (i) for PFC. (colored stars for cre^+^, gray stars for cre^-^, * p < 0.05, ** p< 0.01, *** p < 0.001, Wilcoxon signed-rank test; black stars for comparison cre^+^ vs. cre^-^: * p < 0.05, Wilcoxon rank-sum test). **(C) (i)** Left, imaginary coherence between OB and LEC before (gray) and during (yellow) light stimulation. Right, MI of LFP coherence averaged for different frequency bands for cre^+^ (yellow) and cre^-^ (black) mice. **(ii)** Same as (i) for OB and HP. **(iii)** Same as (i) for OB and PFC. (colored stars for cre^+^: * p < 0.05, ** p < 0.01, *** p < 0.001, Wilcoxon signed-rank test; black stars for comparison cre^+^ vs. cre^-^: * p < 0.05, Wilcoxon rank-sum test).

These results indicate that activation of M/TCs not only induces beta oscillations in OB but also increases the 12-30 Hz oscillatory coupling between OB and downstream cortical areas.

### Inhibition of M/TC output reduces oscillatory power as well as neuronal firing in OB, LEC, and HP

To elucidate whether M/TC activity is necessary for the generation of oscillatory activity in downstream areas, in a first set of experiments, we used inhibitory DREADDs (hM4D(Gi)) that block vesicle release when expressed in M/TCs by cre-dependent virus vector injection (AAV9-EF1a-DIO-hM4D(Gi)-mCherry) at P1 (Fig 6A). At P8, M/TC soma as well as their axons forming the lateral olfactory tract (LOT), which targets the posterior part of the cerebrum, expressed hM4D(Gi)-mCherry (Fig 6B). We performed extracellular recordings of LFP and SUA from OB, LEC, and HP of P8-10 mice (n=35) before (baseline, 20 min) and after (40 min) subcutaneous injection of C21 (3 mg/kg), a synthetic activator of DREADDs [52] (Fig 6A). Since the impact of OB activation on PFC was rather weak, we did not monitor its activity during OB silencing.

**Fig 6.**
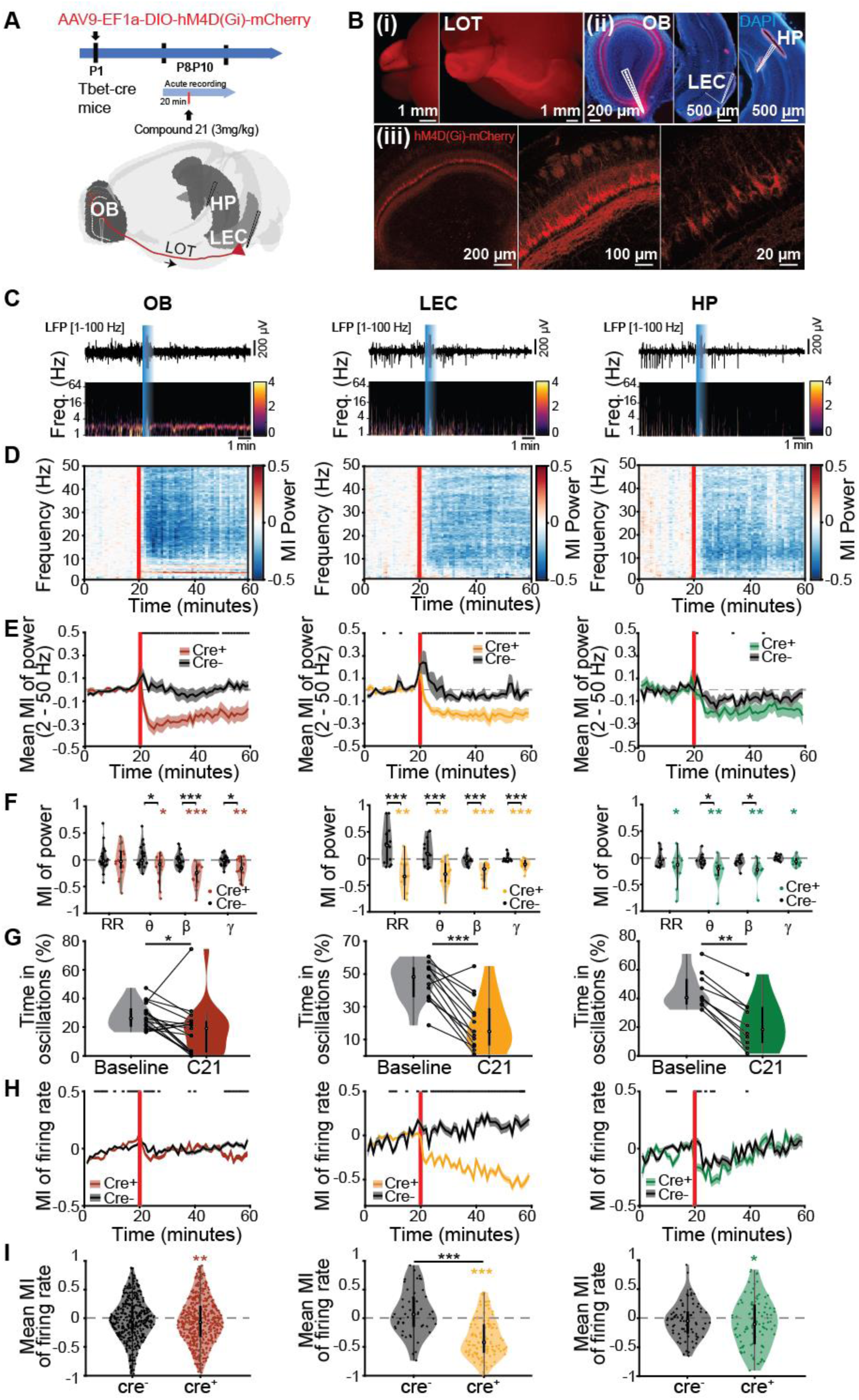
Effects of silencing M/TC output by inhibitory DREADDs on the oscillatory activity in OB, LEC, and HP. **(A)** Top, schematic of the experimental protocol. Bottom, schematic of recording configuration for simultaneous extracellular recordings in OB, LEC, and HP (Brainrender: [48]). **(B) (i)** Photograph of the dorsal (left) and ventral side (right) of a brain from a P8 Tbet-cre^+^ mouse showing mCherry (red) expression in the OB and M/TC axonal projections (LOT) to PIR and LEC. **(ii)** Digital photomontages displaying the DiI labeled electrode track (red) in DAPI (blue) stained slices including the OB (left), LEC (middle), and HP (right) from a P10 mouse. (iii) Confocal images displaying the MCL of the right OB at different magnifications. MC bodies, as well as dendrites, express mCherry. **(C)** Representative LFP traces recorded in the OB, LEC, and HP during C21 injection accompanied by the corresponding wavelet spectra. **(D)** Color-coded MI of LFP power before and after C21 injection in OB (left), LEC (middle), and HP (right). Vertical red lines correspond to the C21 injection. **(E)** Plots displaying the MI of LFP power averaged for 2 to 50 Hz before and after C21 injection in cre^+^ (colored) and cre^-^ (black) mice for OB (right, red), LEC (middle, yellow), and HP (right, green). Vertical red lines correspond to the C21 injection. (black line: p < 0.05, Wilcoxon rank-sum test). **(F)** MI of LFP power averaged for different frequency bands for cre^+^ (colored) and cre^-^ (black) mice for OB (left), LEC (middle), and HP (right). (Wilcoxon signed-rank test, colored stars for cre^+^: * p < 0.05, ** p < 0.01, *** p < 0.001; black stars for comparison cre^+^ vs. cre^-^: * p < 0.05, ** p < 0.01, *** p < 0.001, Wilcoxon rank-sum test). **(G)** Violin plots displaying the percentage of time spend in discontinuous oscillatory events before (baseline, gray) and after C21 injection (C21, colored). Black dots and lines correspond to individual animals. (* p < 0.05, ** p < 0.01, *** p < 0.001, Wilcoxon signed-rank test). **(H)** Line plots displaying the MI of averaged SUA firing rates before and after C21 injection in cre^+^ (colored) and cre^-^ (black) mice for OB (left, red), LEC (middle, yellow), and HP (right, green). Vertical red lines correspond to the C21 injection. (black line: p < 0.05, Wilcoxon signed-rank test). **(I)** Violin plots displaying the MI of averaged SUA firing rates after C21 injection for cre^-^ (black) and cre^+^ mice (colored) recorded in OB (left, red), LEC (middle, yellow), and HP (right, green). Red and black dots correspond to individual units. (colored stars for cre^+^: * p < 0.05, ** p < 0.01, *** p < 0.001, Wilcoxon signed-rank test; black stars for comparison cre^+^ vs. cre^-^: *** p < 0.001, Wilcoxon rank-sum test).

C21 caused broadband power reduction in OB that reached a maximum magnitude within 5 min after the injection (Figs 6C, 6D, 6E, 6F, and S9 Table) and persisted for at least 2 h (S5C Fig). The occurrence of discontinuous oscillatory events was lower after C21 injection in OB (Fig 6G and S10 Table), indicating that M/TC activity is involved in the generation of discontinuous events in OB. Solely, the continuous RR in OB was not affected by the activation of DREADDs (Fig 6F and S9 Table). Moreover, silencing the M/TC output led to a broadband reduction of oscillatory power in LEC and HP (Figs 6C F and Table S9). Correspondingly, the time spend in oscillatory events in LEC and HP decreased after inhibition of M/TC output (Fig 6G and S10 Table). In contrast, for cre^-^ mice LFP power and time spent in oscillatory events did not differ before and after C21 injection (S5A and S5B Figs, S9 and S10 Tables).

Next, we monitored the effects of chemogenetic silencing of M/TCs on the neuronal firing of downstream areas. Inhibitory DREADDs have been described to mainly reduce the vesicle release in the expressing neurons, while having little, if any, impact on their ability to generate action potentials [53,54]. Indeed, C21 injection had a weak effect on SUA in OB (cre^+^: med MI: -0.071, iqr: -0.322 – 0.208, n=512, p=0.003, Wilcoxon signed-rank test; cre^-^: med MI: -0.028, iqr: -0.259 – 0.207, n=418, p=0.171, Wilcoxon signed-rank test; cre^+^ vs. cre^-^: p=0.198, Wilcoxon rank-sum test) (Figs 6H and 6I). In particular, the neuronal firing within the first 10 min after C21 injection decreased (Fig 6H), being most likely the result of weaker network interactions within the OB. The DREADDs manipulation affected not only the network and neuronal activity in OB but also the spike timing by oscillations. In line with the results of spike-triggered power (STP) analysis, C21 injection decreased the ability of SUA to entrain the OB in theta, beta, and gamma rhythms (S6A Fig and S11 Table). STP for RR was comparable in the presence and absence of C21 (S11 Table). The temporal relationship between OB spikes and oscillatory events in OB was also assessed by calculating the phase-locking of SUA to RR and beta rhythm, respectively. In line with the results of STP analysis, the phase-locking to beta was reduced after C21 injection for all units (baseline: med: 0.105, iqr: 0.061 – 0.152; C21: med: 0.093, iqr: 0.053 – 0.139; n_units_=524 from 16 mice, p=0.003, LMEM) as well as for units that were significantly locked during the baseline period (baseline: med: 0.120, iqr: 0.084 – 0.165 C21: med: 0.101, iqr: 0.065 – 0.148; n_units_=273 from 16 mice, p=7.15*10^-5^, LMEM). In contrast, C21 had no effect on the number of significantly locked units (baseline: 52.1%, C21: 51.9%, p=1, Fisher’s exact test) (S6C Fig). The phase coupling to RR (baseline: med: 0.094, iqr: 0.051 – 0.153; C21: med: 0.159, iqr: 0.078 – 0.310; n_units_=524 from 16 mice, p=2.20*10^-16^, LMEM) (S6B Fig) and the fraction of significantly locked units to the RR (baseline: 46.0%, C21: 71.9%, p=1.42 *10^-17^, Fisher’s exact test) were increased after C21 injection.

Silencing the M/TC output strongly reduced the LEC firing (cre^+^: med MI: -0.420, iqr: - 0.598 – -0.108, n=126, p=2.96*10^-16^, Wilcoxon signed-rank test; cre^-^: med MI: 0.069, iqr: - 0.144 – 0.364, n=49, p=0.168, Wilcoxon signed-rank test; cre^+^ vs. cre^-^: p=3.87*10^-10^, Wilcoxon rank-sum test), the effects lasting > 1 hour after C21 injection (Figs 6H and 6I). In contrast, silencing of M/TC output had a shorter (∼20 min) and weaker impact on hippocampal firing (cre^+^: med MI: -0.102, iqr: -0.438 – 0.230, n=102, p=0.036, Wilcoxon signed-rank test; cre^-^: med MI: -0.047, iqr: -0.256 – 0.109, n=74, p=0.119, Wilcoxon signed-rank test; cre^+^ vs. cre^-^: p=0.484, Wilcoxon rank-sum test).

Since the impact of DREADD-induced silencing of OB synaptic outputs on the LEC and HP activity results from both mono and polysynaptic connectivity, in a second set of experiments, we dissected the role of the direct monosynaptic OB-to-LEC pathway using recently developed inhibitory opsins. We expressed the targeting-enhanced mosquito homolog of the vertebrate encephalopsin (eOPN3) that selectively suppresses neurotransmitter release at presynaptic terminals [55], into M/TCs (Fig 7A). Light stimulation (473 nm) of eOPN3-expressing MC terminals in LEC (Fig 7B) reduced the power of entorhinal beta band activity (med: -0.213, iqr: -0.269 – -0.095, p = 0.008, n=9, Wilcoxon signed-rank test), yet not of RR (med: -0.109, iqr: -0.252 – 0.079, p = 0.496, n=9, Wilcoxon signed-rank test) as well as theta band (med: -0.156, iqr: -0.231 – 0.011, p = 0.098, n=9, Wilcoxon signed-rank test) and gamma band oscillations (med: -0.048, iqr: -0.107 – -0.022, p = 0.055, n=9, Wilcoxon signed-rank test) in LEC (Figs 7C and 7Dii). Consequently, the optogenetic silencing led to a reduction of oscillatory power in HP and PFC in a frequency range spanning from RR to beta (HP: RR: med: -0.122, iqr: -0.266 – -0.095, p = 0.016; theta: med: -0.242, iqr: -0.363 – -0.122, p = 0.031; beta: med: -0.282; iqr: -0.345 – -0.131, p = 0.016; gamma: med: -0.073, iqr: -0.136 – 0.011, p = 0.078; n=7; PFC: RR: med: -0.181, iqr: -0.375 – -0.072, p = 0.016; theta: med: -0.166, iqr: 0.361 – -0.080, p = 0.047; beta: med: -0.132; iqr: -0.208 – -0.044, p = 0.047; gamma: med: - 0.054, iqr: -0.092 – 0.011, p = 0.219; n=7; Wilcoxon signed-rank test) (Figs 7C, 7Diii and 7Div). Of note, beta and gamma power in OB was reduced following the silencing of MC terminals in LEC (RR: med: -0.002, iqr: -0.099 – 0.086, p = 0.820; theta: med: -0.039, iqr: -0.127 – 0.034, p = 0.426; beta: med: -0.139, iqr: -0.253 – -0.074, p = 0.020; gamma: med: -0.062, iqr: -0.111 – -0.020, p = 0.020; n=9; Wilcoxon signed-rank test) (Figs 7C and 7Di), most likely as result of reduced centrifugal feedback from LEC and HP [56].

**Fig. 7:**
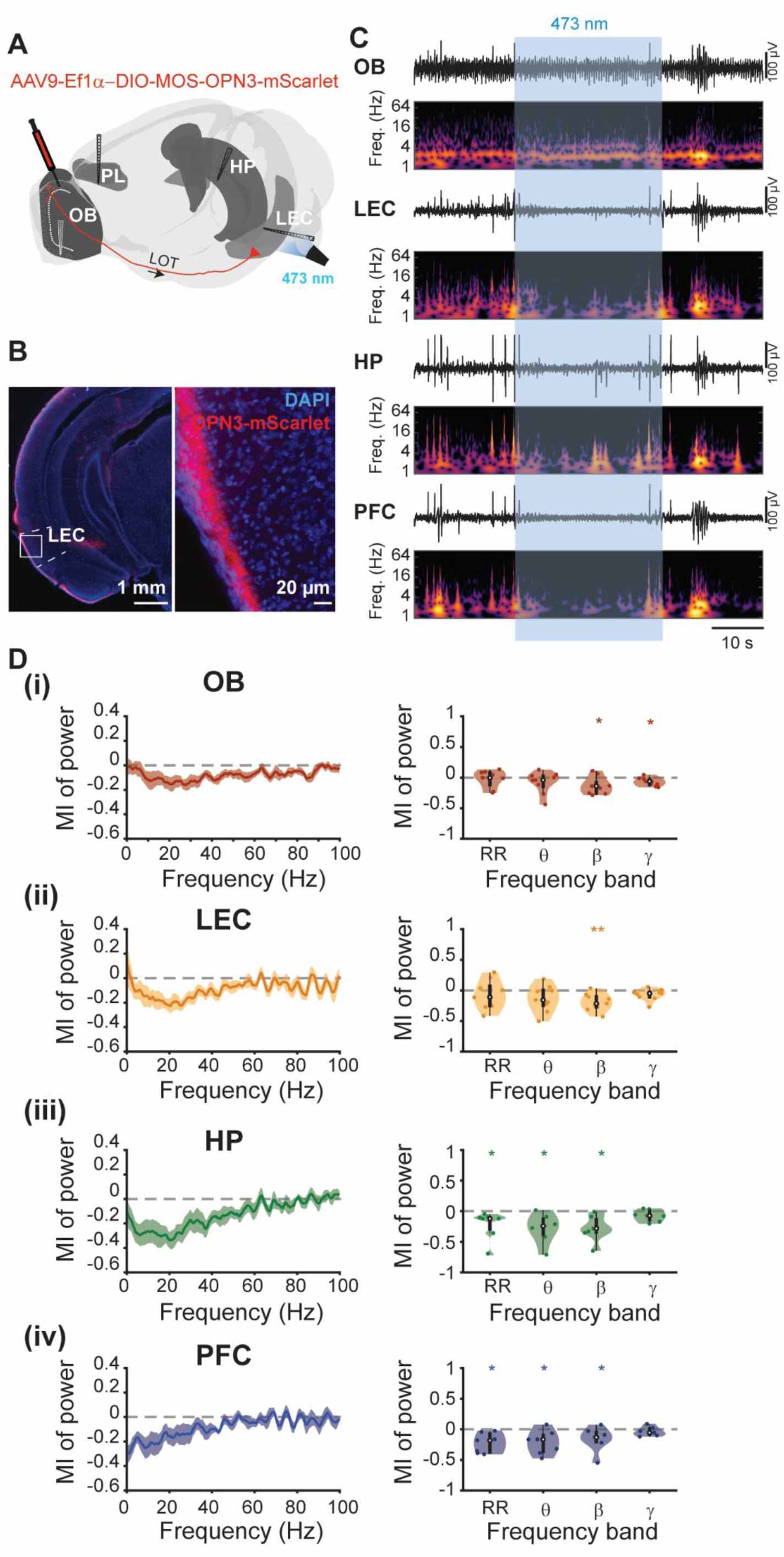
Light-induced inhibition of MC axonal terminals in LEC. **(A)** Top, schematic of recording configuration for simultaneous extracellular recordings in OB, LEC, HP, and PFC (Brainrender: [48]). **(B)** Photograph of a DAPI (blue) stained slice from a P8 Tbet-cre^+^ mouse showing mScarlet (red) expression in MC axonal projections (LOT) to LEC in two different magnifications. **(C)** Representative LFP traces recorded in the OB, LEC, HP, and PFC during stimulation of OPN3-transfected M/TCs accompanied by the corresponding wavelet spectra. Vertical black lines correspond to 100 µV. **(D) (i)** Left, plot of MI for OB power during optogenetic activation of OPN3 in MC axon terminals (red). Right, MI of LFP power averaged for different frequency bands for OB. **(ii)** Same as (i) for LEC (yellow). **(iii)** Same as (i) for HP (green). **(iv)** Same as (i) for PFC (blue). (* p < 0.05, ** p < 0.01, *** p < 0.001, Wilcoxon signed-rank test).

These results indicate that silencing the M/TC output decouples neuronal firing from beta oscillations in OB and decreases the oscillatory power and neuronal firing in LEC, as a first downstream station of OB projections. On its turn, the weaker drive from LEC leads to poor oscillatory entrainment of HP, yet without significant change in its neuronal firing.

### Inhibition of M/TC output reduces the communication between OB and downstream cortical areas

To back up the hypothesis that the M/TC activity controls the developmental entrainment of limbic circuits, we monitored the communication between OB and downstream areas during the silencing of M/TC output with DREADDs by using three distinct measures. First, we assessed the synchrony between OB, LEC, and HP by calculating the imaginary coherence in different frequency bands before (baseline) and after C21 injection (C21) (Figs 8A and 8B). MIs for beta coherence between OB and LEC, and OB and HP were significantly reduced after C21 injection. In contrast, the coherence in other frequency bands was not affected by C21 injection (Figs 8A and 8B, S12 Table). Moreover, the C21-induced changes in the beta band were not detected in cre^-^ mice (S7A Fig and S12 Table).

**Fig 8.**
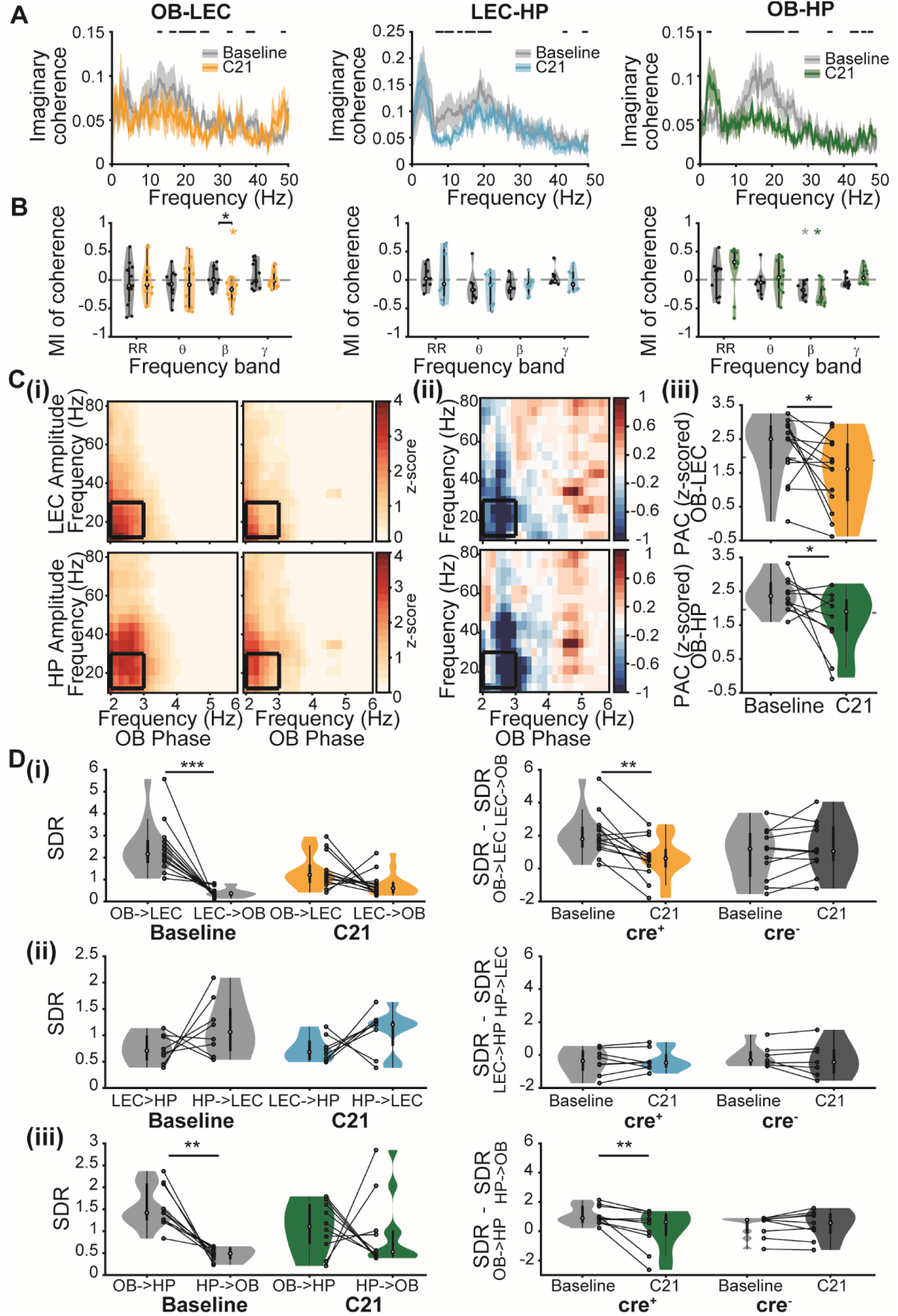
Modulation of functional communication within olfactory-cortical networks through silencing the M/TC output by inhibitory DREADDs. **(A)** Imaginary coherence calculated for OB-LEC (left, yellow), LEC-HP (middle, light blue), and OB-HP (right, green), before (baseline, gray) and after C21 injection (C21, colored). (black line: p < 0.05, Wilcoxon rank-sum test). **(B)** MI of coherence averaged for different frequency bands between OB and LEC (left, yellow), LEC and HP (middle, light blue), and OB and HP (right, green), for cre^+^ (colored) and cre^-^ (black) mice. (colored stars for cre^+^, gray stars for cre^-^: * p < 0.05, Wilcoxon signed-rank test; black stars for comparison cre^+^ vs. cre^-^: * p < 0.05, Wilcoxon rank-sum test). **(C) (i)** Z-scored phase-amplitude coupling (PAC) between OB phase and LEC (top) and HP (bottom) amplitude, before (baseline) and after C21 injection (C21). **(ii)** Difference of PAC values after and before C21 injection for OB-LEC (top) and OB-HP (bottom). **(iii)** PAC averaged for RR-beta coupling (black box in (i)) for OB-LEC (top) and OB-HP (bottom), before (baseline, gray) and after C21 injection (colored). Dotted gray line corresponds to a z-score of 1.96. (* p < 0.05, Wilcoxon signed-rank test). **(D) (i)** SDR calculated for OB and LEC. Left, SDR values for OB -> LEC and LEC -> OB before (baseline, gray) and after C21 injection (C21, yellow). Right, difference of SDR values for both directions for cre^+^ and cre^-^ mice. **(ii)** Same as (i) for LEC and HP (blue). **(iii)** Same as (i) for OB and HP (green). Black dots and lines correspond to individual animals. (** p < 0.01, *** p < 0.001, Wilcoxon signed-rank test).

Second, we calculated the phase-amplitude coupling (PAC) to elucidate the role of M/TCs in the modulation of cortical beta oscillations by the RR phase in OB. C21 injection significantly reduced the z-scored PAC values between the OB RR phase and the amplitude of beta oscillations in LEC (baseline: med: 2.499, iqr: 1.624 – 2.883; C21: med: 1.608, iqr: 0.674 – 2.361, n=13, p=0.017, Wilcoxon signed-rank test) and HP (baseline: med: 2.363, iqr: 2.135 – 2.764; C21: med: 1.907, iqr: 1.319 – 2.248, n=10, p=0.037, Wilcoxon signed-rank test) (Fig 8C). Additionally, fewer mice showed significant RR-beta PAC values after C21 injection (z-score > 1.96) in LEC (baseline: 53.9 % vs. C21: 39.8 %) and HP (90 % vs. 50 %).

Third, we tested the effect of C21 on the directionality of interactions between OB, LEC, and HP (Fig 8D). We calculate the SDR and found that the prominent drive from OB to LEC was absent after silencing of M/TC output, the values for OB → LEC and LEC → OB being comparable (Fig 8Di, S13, and S14 Tables). Similarly, the drive from OB to HP was disrupted by C21 injection (Fig 8Diii, S13 and S14 Tables). In contrast, the directionality of interactions between LEC and HP was not affected by C21 injection. As reported for the baseline conditions, the mutual interactions LEC-HP persisted after M/TC silencing (Fig 8Dii, S13, and S14 Tables). Moreover, the C21-induced changes in directionality were not detected in cre^-^ mice (S7B Fig, S13 and S14 Tables).

Thus, these results show that the M/TC activity is critical for the communication between OB and its downstream cortical areas.

## DISCUSSION

Long-range interactions within limbic circuits emerge early in life [57], yet it is still unknown whether the coordinated activity patterns underlying the coupling are endogenously generated or emerge as result of the driving force of sensory systems. Several distinct activity patterns have been identified in developing cortical-hippocampal networks. Apart from discontinuous theta and beta oscillations synchronizing LEC-HP-PFC networks at neonatal age [21,58,59], spontaneous twitches during active sleep drive early sharp waves in the developing HP [15,60]. Besides muscle twitches [14,61] and passive tactile sensation, olfactory inputs are likely candidates for the instruction of limbic circuitry development during the first two postnatal weeks. Newborn rodents are not only able to smell from birth on but, importantly, also use olfactory information for learning and cue-directed behaviors such as localization of the nipples of the dam [29,62]. A first piece of evidence for the critical role of olfaction for limbic development is the fact that the neonatal OB shows functional coupling with the LEC, the gatekeeper of the limbic circuitry, during discontinuous network oscillations in the theta-beta frequency range as well as in the continuous respiration-related rhythm (RR) in anesthetized mice [43]. Here, we extended these findings and uncovered that MC firing sets a beta band entrainment also in downstream areas, such as HP and PFC in awake neonatal mice (P8-10). The temporal dynamics of oscillatory and firing activity revealed that in periods with and without active odor sampling, OB drives the activation of limbic circuits. Layer-specific analysis of SUA revealed that M/TC activation leads to a complex entrainment of the OB microcircuit that results in an augmented firing rate also for interneurons in the GCL, EPL, and GL. Experimental and modeling studies have shown that both beta and gamma oscillations in the OB rely on dendro-dendritic interactions between M/TCs and GCs [33,50,61–65]. In adults, the emergence of gamma and beta oscillations is controlled by different excitability states of GCs as well as their dependency on centrifugal input, with beta oscillations relying on a higher GC excitability and centrifugal feedback projections [40,63,64]. However, gamma oscillations are absent in the neonatal OB, most likely as a result of the late functional integration of interneurons into local circuits and the different biophysical properties of MCs and GCs during development [66-68]. Instead, discontinuous beta band oscillations are present not only in OB but also in other sensory and limbic areas [19]. In the neonatal PFC, they have been shown to accelerate along development until reaching the gamma-band range at juvenile age [69]. Similarly, acceleration of beta to gamma oscillations takes place in V1 during the critical period for vision [70,71]. Whether the beta band activity in OB undergoes a similar transition to faster rhythms and how this process is controlled by interactions within OB and by feedback projections from PiR and LEC remain to be elucidated.

In a previous study, we have shown that in anesthetized mice, OB-driven entrainment of the neonatal LEC is strongest in the theta frequency range [43]. Here we show that in awake mice, the MC-driven synchronization of OB and LEC shifts to beta frequency range. Urethane anesthesia has been shown to decrease beta and gamma oscillations in the OB of adult rats and dampens spontaneous and odor-evoked GC responses while enhancing MC firing in response to odors [72-75]. Thus, the anesthesia-induced reduction in fast frequency oscillations might be due to decreased dendro-dendritic inhibition on MCs in adults [74]. As GCs are strongly targeted by centrifugal projections and beta oscillations in OB are dependent on centrifugal inputs [76], the absent beta synchronization between OB and downstream cortical brain areas, which we have observed during anesthesia [43], might be due to reduced centrifugal input to the OB [75].

The present data show that the OB network activation entrains downstream cortical areas in beta oscillations in neonatal awake mice (P8-10). In adult rodents, the axonal terminals of MCs have been found to target fan and pyramidal neurons in LII/III of LEC that, on their turn, relay this information to the HP [77,78]. The axonal projections from layer II/III LEC pyramidal neurons to CA1 are involved in associative odor learning in adults [79]. Already at neonatal age, MC axons reach layer I of LEC [43,80]. Here, projections of layer II/III neurons that target the HP were detected and they might establish synaptic contacts with the MC axons. Optogenetic stimulation revealed that the activation of M/TCs induced delayed firing of LEC neurons and HP neurons, indicating that the pathway OB-to-HP is indeed already functional from birth on. CA1 receives entorhinal input not only via the direct performant path but also through the tri-synaptic path, spanning DG and CA3 [81]. The long latency (∼ 60 ms) in light-induced CA1 firing might, therefore, be partly mediated by the tri-synaptic path. Further, LEC receives not only inputs directly from OB but also indirectly via PiR [82]. While the rather long and broad activity increase in LEC after MC stimulation might argue for PiR involvement, a prominent monosynaptic drive from OB to LEC is present too and controls the downstream HP, as demonstrated by the OPN3-induced silencing. Of note, it was recently shown that a distinct but rather small population of LEC layer II neurons directly projects to the neonatal PFC [26], yet light stimulation of M/TCs did not recruit it.

Coordinated activity patterns in OB organized by MCs promote not only neuronal firing but also network activation in downstream areas. Olfactory stimulation with different odors leads to an immediate increase of firing in the OB as well as a broadband increase in power in OB, LEC, HP, and PFC. Similarly, ramp light stimulation of M/TCs led to an increase in beta band power in LEC, HP, and PFC. This power surge was accompanied by increased long-lasting SUA firing in all three brain areas, indicating that the initial activation of neurons is followed by activation of the local networks in LEC, HP, and PFC. This long-lasting local network activation could be caused by engaging the entorhinal-hippocampal loop, leading to a reactivation of LEC caused by HP output. Conversely, blocking vesicle release on MC synapses by DREADDs reduced the broadband power as well as neuronal firing in LEC and HP. Selective silencing of the MC axons terminating in LEC using eOPN3 confirmed that the direct projections from OB to LEC are important for the entrainment of oscillatory activity in LEC, HP, and PFC. Moreover, coherence analysis revealed increased oscillatory, mainly beta band coupling, between OB and cortical areas during olfactory and ramp stimulation, whereas inhibition of M/TCs vesicle release reduced the drive OB → LEC and OB → HP as well as RR-beta cross-frequency coupling and beta coherence between OB-LEC and OB-HP. Considering that the artificial activation of MCs leads to a similar entrainment of downstream areas as observed during olfactory sampling, these results identify the MC-driven beta rhythm as a potential mechanism of long-range communication between OB and downstream cortical networks.

What might be the relevance of OB-controlled beta band activation of cortical circuits during early postnatal development? Beta oscillations have been reported to play a key role in working memory and decision-making in adult humans [83]. Further, prominent beta band synchrony between cortical areas has been identified during olfactory-guided memory and decision-making tasks in rodents [38,40,42,84]. A similar, but sniffing-independent increase in hippocampal beta oscillations has been observed during an object learning task [85]. Moreover, the firing of beta-entrained CA1 interneurons during an odor-place associative memory and decision-making task is related to an accurate performance, indicating that beta oscillations enable temporal coordination and recruitment of neurons within functional cell assemblies [42,84]. In line with these experimental data, modeling confirmed that beta oscillations optimally contribute to the coupling of cell assemblies over long axonal conductance delays [86-88]. During development, discontinuous beta band events that have been identified in PFC, HP, and LEC might facilitate the formation of initial cell assemblies with relevance for cognitive performance later in life. Moreover, sensory feedback signals evoked by myoclonic twitches synchronize the primary sensory cortex and the HP in the beta frequency range during sleep [14]. We previously showed that interfering with beta band oscillations during a defined developmental period causes network miswiring and poor behavioral performance in adult mice [28]. Similarly, in a mouse model of psychiatric risk reduced beta band activity at neonatal age has been found to correlate with later cognitive deficits [22,26]. Moreover, reduced rates of hippocampal beta oscillations were observed in an animal model of epilepsy [89]. Here, we identified the olfactory activity as a prominent driver of these early beta oscillations. Thus, olfactory sensory inputs, possibly in conjunction with other sensory signals such as reafferent signaling following spontaneous twitches, could be important for the synchronization of limbic network activity during the second postnatal week. The results let us hypothesize that transient disturbance of neonatal olfactory processing precludes the functional refinement of entorhinal-hippocampal-prefrontal circuits, ultimately leading to cognitive deficits in adulthood. Further research is warranted to directly test this hypothesis and elucidate the role of early activity patterns in OB for cognitive development.

## MATERIALS AND METHODS

### Ethical Approval

All experiments were performed in compliance with the German laws and the guidelines of the European Union for the use of animals in research (European Union Directive 2010/63/EU) and were approved by the local ethical committee (Behörde für Gesundheit und Verbraucherschutz Hamburg, ID 15/17).

### Animals

Time-pregnant C57Bl/6/J and Tbet-cre mice from the animal facility of the University Medical Center Hamburg-Eppendorf were housed individually in breeding cages at a 12h light / 12h dark cycle and fed ad libitum. Offspring (both sexes) where injected with either AAV9-Ef1a-DIO-hChR2(E123T_T159C)-EYFP (Addgene, Plasmid #35509), AAV9-EF1a-DIO-hM4D(Gi)-mCherry (Addgene, Plasmid #50461) or AAV9-EF1a-DIO-MOS-OPN3-mScarlet virus at postnatal day (P) 0 or 1. Genotypes were determined using genomic DNA and following primer sequences (Metabion, Planegg/Steinkirchen, Germany) as described previously [43]: for Cre: PCR forward primer 5′-ATCCGAAAAGAAAACGTTGA-3′ and reverse primer 5′-ATCCAGGTTACGGATATAGT-3′. The PCR reactions were as follows: 10 min at 95 °C, 30 cycles of 45 s at 95 °C, 90 s at 54 °C, and 90 s at 72 °C, followed by a final extension step of 10 min at 72 °C. In addition to genotyping, EGFP expression in OB was detected using a dual fluorescent protein flashlight (Electron microscopy sciences, Hatfield, PA, USA) prior to surgery. At P8-10 cre^-^ and cre^+^ mice underwent light stimulation or Compound 21 injections and *in vivo* multi-side electrophysiological recordings.

### Surgical procedures and recordings

#### Virus injection for transfection of MTCs with ChR2 and hm4D(Gi)

For transfection of M/TCs with the ChR2 derivate E123T/T159C, eOPN3 or inhibitory DREADDs (hm4D(Gi)), P0-1 pups were fixed into a stereotaxic apparatus and received unilateral injections of one of two viral constructs (AAV9-Ef1a-DIO hChR2(E123T/T159C)-EYFP, 200 µl at titer ≥ 1×10¹³ vg/mL, Plasmid, #35509, Addgene, MA, USA; AAV9-EF1a-DIO-hM4D(Gi)-mCherry, 200 µl at titer ≥ 1×10^14^ vg/mL Plasmid #50461, Addgene, MA, USA; AAV9-EF1a-DIO-MOS-OPN3-mScarlet, 200 µl at titer 8.7 ×10^11^ vg/ml, Plasmid #131002, Addgene, MA, USA). The virus was produced by Addgene or the Virus Facility of the University Medical Center Eppendorf. A total volume of 200 nl was slowly (200 nl/min) delivered at a depth of around 0.5 mm into the right OB using a micropump (Micro4, WPI, Sarasota, FL). Following injection, the syringe was left in place for at least 30 s to avoid reflux of fluid. Pups were maintained on a heating blanket until full recovery and returned to the dam.

#### Virus injection for tracing

For the transfection of M/TC axons with EYFP and the retrograde labeling of HP-projecting neurons with mCherry, P0-1 pups received the viral construct AAV9-hSyn-hChR2(H134R)-EYFP (200 µl at titer ≥ 1×10¹³ vg/mL, #26973-AAV9, Addgene, MA, USA) into the OB and the retrograde virus AAVrg-CamKIIα-mCherry (80 µl at titer ≥ 7×10¹² vg/mL, #114469-AAVrg, Addgene, MA, USA) into the HP. Virus injection was performed similarly as for the transfection of M/TCs with ChR2 or hm4D(Gi). After 10 days, the brains of investigated mice were perfused with 4% paraformaldehyde (PFA), sliced and MC axons and HP-projecting neurons in LEC and PIR were imaged using a confocal microscope.

#### Surgical procedure for electrophysiology

For in vivo recordings, P8-10 mice underwent surgery according to previously described protocols [20,43,44]. Under isoflurane anesthesia (induction: 5 %, maintenance: 2.5 %, Forane, Abbott), the skin above the skull was removed and 0.5 % bupivacaine / 1 % lidocaine was locally applied on the neck muscles. Two plastic bars were mounted on the nasal and occipital bones with dental cement. The bone above the right OB (0.5-0.8 mm anterior to frontonasal suture, 0.5 mm lateral to inter-nasal suture), LEC (0 mm posterior to lambda, 6-7.5 mm lateral from the midline), HP (2.5 mm anterior to lambda, 3.5 mm lateral from the midline) and PFC (0.5 mm anterior to bregma, 0.1-0.5 mm lateral from the midline) was carefully removed by drilling a hole of < 0.5 mm in diameter. Throughout surgery and recording session the mice were maintained on a heating blanket at 37°C.

#### Multi-site electrophysiological recordings in vivo

Three-side or four-side recordings were performed in non-anesthetized P8-10 mice. For this, one-shank electrodes (NeuroNexus, MI, USA) with 16 recording sites (0.4-0.8 MΩ impedance, 50 µm inter-site spacing for recordings in OB and HP, 100 µm inter-site spacing for recordings in LEC and PFC) were inserted into OB (0.5-1.8 mm, angle 0°), LEC (for 4-side recordings, depth: 2 mm, angle: 180°; for 3-side recordings, depth: 2-2.5 mm, angle: 10°), HP (1.3-1.9 mm, angle 20°) and PFC (1.8-2.1 mm, angle 0°). For light stimulation one-shank optrodes (NeuroNexus, MI, USA) with the same configuration as the electrodes were inserted in the OB. Before insertion, the electrodes were covered with DiI (1,1’-Dioctadecyl-3,3,3’,3’-tetramethylindocarbocyanine perchlorate, Molecular Probes, Eugene, OR). A silver wire was inserted into the cerebellum and served as a ground and reference electrode. Before data acquisition, a recovery period of 20 min following the insertion of electrodes was provided. Extracellular signals were band-pass filtered (0.1 Hz-9 kHz) and digitized (32 kHz or 32,556 kHz) by a multichannel amplifier (Digital Lynx SX; Neuralynx, Bozeman, MO; USA) and Cheetah acquisition software (Neuralynx). Spontaneous activity was recorded for at least 20 min before light stimulation or Compound 21 (C21, Hellobio, Ireland) injection. The position of recording electrodes in OB, LEC, HP, and PFC was confirmed after histological assessment *post-mortem*. For the analysis of LFP in OB, the recording site centered in the EPL was used, whereas for HP the recording site located in the CA1 was considered. For analysis of LFP in LEC only recording sites that were histologically confirmed to be located in superficial entorhinal layers were used. Similarly, only recordings sites confined to the prelimbic sub-division of PFC were considered. For the analysis of spiking activity, all recording sites confirmed to be located in the areas of interest (OB, LEC, HP, and PFC) were considered. When necessary, spikes recorded in OB were assigned to specific layers according to the location of recording sites.

#### Morphology

Mice were anesthetized with 10% ketamine (Ketamidor, Richter Pharma AG, Germany) / 2% xylazine (Rompun, Bayer, Germany) in 0.9% NaCl solution (10 µg/g body weight, i.p.) and transcardially perfused with Histofix (Carl Roth, Germany) containing 4% PFA. Brains were postfixed in 4% PFA for 24 h and sliced. Slices (100 µm-thick) were mounted with Fluoromount containing DAPI (Sigma-Aldrich, MI, USA). The positions of the DiI-labeled extracellular electrodes in the OB, LEC, HP, and PFC were reconstructed to confirm their location. Virus expression was verified by EYFP (for ChR2) or mCherry (for hM4D(Gi)) fluorescence in the right OB. For confocal imaging of EYFP or mCherry fluorescence in M/TCs, HP, and LEC, 50 µm-thick slices mounted with Vectashield (CA, USA) were used.

#### Light stimulation

Activation of M/TCs was achieved by either ramp or pulse light stimulation applied using a diode laser (473 nm; Omicron, Austria) which was controlled by an arduino uno (Arduino, Italy). For ramp stimulation, a light stimulus with linear increasing power (3 s rise time) was presented 30-60 times. For pulse stimulation 3 ms light pulses at 2 Hz were delivered. Laser power was adjusted for every recording (1.37-5.15 mW) to reliably induce neuronal firing. For the inactivation of neurotransmitter release from axon terminals in LEC light stimulation (10 s) was applied using a diode laser (473 nm; Omicron, Austria).

#### Olfactory stimulation

For olfactory stimulation a custom-made olfactometer was used and odors were applied for 2 s triggered by respiration and controlled by an arduino (Arduino, Italy). Two to four different odors ((R)-(+) Limonene [Alfa Aesar], 1-Octanol [Acros Organics], Vanillin, and Isoamyl acetate [Sigma-Aldrich], 1% in mineral oil [Sigma-Aldrich]) were delivered in a randomized order using mass flow controllers.

#### Compound 21 injection

Compound 21 (3 mg/kg solved in 0.9% NaCl) was injected subcutaneously after >20 min recording of baseline activity, while the mouse was fixed in the stereotaxic apparatus. The activity was recorded for 40-120 min post-injection.

### Data Analysis

#### LFP analysis

Data were analyzed offline using custom-written scripts in the MATLAB environment (MathWorks, Natick, MA). Data were first low-passed filtered (<100 Hz) using a third-order Butterworth filter before down-sampling by factor 20 to 1.6 kHz to analyze LFP. All filtering procedures were performed in a manner preserving phase information.

#### Detection of oscillatory activity

Discontinuous network oscillations in the LFP recorded from OB, LEC, and HP before and after C21 injection were detected using a previously developed unsupervised algorithm [90]. Briefly, deflections of the root mean square of band-pass filtered (4-100 Hz) signals exceeding a variance-depending threshold (2 times the standard deviation from the mean) were assigned as oscillatory periods. Only oscillatory periods lasting at least 1 s were considered for analysis.

#### Power spectral density

Power spectral density was analyzed for either the entire baseline period, 2 s long periods before (Pre), and during light ramp stimulation (Stim) for recordings combined with optogenetic manipulation. For recordings paired with DREADD manipulation, the power was either calculated for every minute or averaged for the entire baseline period (19 min) and post C21 injection period (30 min). Power was calculated using Welch’s method with non-overlapping windows of 2 s (ramp periods) or 3 s length and spectra were smoothened with a moving average of 20 samples. Time-frequency plots of power were calculated with a continuous wavelet transform (Morlet wavelet).

#### Coherence

The imaginary part of coherence, which is insensitive to volume-conduction-based effects [501], was calculated for the same time periods as the power by taking the absolute value of the imaginary component of the normalized cross-spectrum:

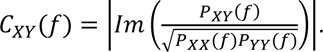

#### Spectral Dependency Ratio

The Spectral Dependency Ratio (SDR) was calculated according to Shajarisales et al. [47] from the power spectral densities (Sx(f) and Sy(f)) of the signals X and Y:

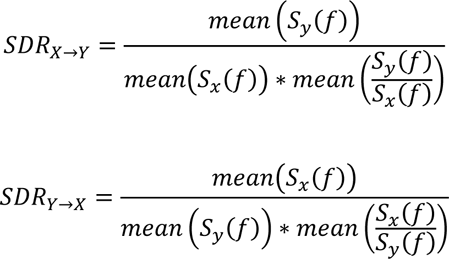

The most likely direction of causation is the one having significantly larger SDR values. (https://github.com/OpatzLab/HanganuOpatzToolbox/tree/master/LFPanalysis/getSDR.m)

#### Spiking analysis

Single units were automatically detected and clustered using the python-based software klusta [91] and manually curated using phy (https://github.com/cortex-lab/phy). The firing rate was computed by dividing the total number of spikes by the duration of the analyzed time window. To assess the spike probability, histograms of spike count using 1 ms bins were calculated for periods around the light pulse (50 ms before to 150 ms after) and normalized to the number of delivered light pulses. For peristimulus time histograms of odor-evoked firing rates, even trials were sorted relative to the peaks obtained from odd trials. Cross-covariance of spike trains was calculated as described previously [43,45]. Briefly, cross-covariance for two spike trains *N_i_* and *N_j_*, was estimated from the cross-correlation histogram 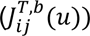 as follows:

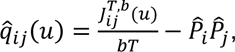

(*b* = binsize, *T* observation period, 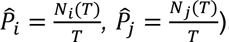. The standardized cross-covariance was calculated as

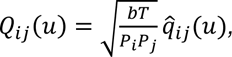

with *P_i_*, *P_j_* being the mean firing rates. Only pairs of units with firing rates > 0.05 Hz and significant standardized cross variance were considered. The Null hypothesis was rejected when 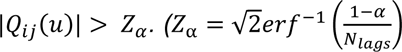; two-tailed critical z value at level *α* = 0.01). The standardized mean cross-variance for one unit was calculated as

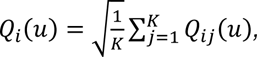

(*K=* number of units in 2. Region) and the mean for all unit pairs as: 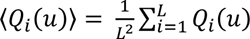.

#### Modulation index

The modulation index (MI) of power, coherence, firing rate, and STP for light stimulation or DREADD manipulation was calculated as

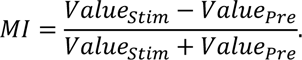

#### Spike-LFP coupling

Phase locking of spiking units to network oscillations was assessed using a previously described algorithm [45]. For this, the LFP signal was bandpass filtered (2-4 Hz (RR), 4-12 Hz (theta), 12-30 Hz (beta), 30-100 Hz (gamma) using a third-order Butterworth filter. The instantaneous phase was extracted using the Hilbert transform on the filtered signal. The coupling between spikes and network oscillations was tested for significance using the Rayleigh test for non-uniformity. For analysis of baseline properties (Fig 1) only neurons that showed significant phase locking were considered for the analysis of the mean phase angle and the locking strength, which was calculated as mean resulting vector length (RVL). For paired comparison of RVLs (Fig 2, S1, S4) all units with a firing rate higher than 0.01 Hz during baseline (18 min) and after C21 injection (18 min, DREADD manipulation) or more than 10 spikes before (Pre) and during (Stim) light ramp pulses were considered. (https://github.com/OpatzLab/HanganuOpatzToolbox/blob/master/Spikes-LFPanalysis/getPPC_PLV.m)

#### Spike-triggered power

Spike-triggered power (STP) was calculated for the same time periods as RVL by taking the mean of the LFP power for 0.4 s long time windows centered on each spike.

#### Phase-amplitude coupling

Phase-amplitude coupling (PAC) between RR phase in OB and beta band amplitude in LEC and HP was calculated as previously described [92]. Briefly, the LFP signals were bandpass filtered and the Hilbert transform was used to extract the phase and amplitude, respectively. Subsequently, the amplitude of the beta-filtered signal in LEC or HP was determined at each phase of the filtered OB signal. The phase was divided into 16 bins and the mean amplitude for each bin was calculated and normalized to the total number of bins. The normalized modulation index (MI) was calculated as the deviation between an empirical and uniform amplitude distribution. MI matrices were z-scored and the average was calculated for RR (2-3 Hz) – beta (12-30 Hz) coupling.

### Statistics

Statistical analysis was performed in MATLAB environment or R Statistical Software. As none of the data sets were normally distributed, data were tested for significance using Wilcoxon rank-sum test (2 unrelated samples) or Wilcoxon sign-rank test (2 related samples). Data (except phase values) are presented as median (med) and interquartile range (iqr). Outlier removal was applied to paired data points if the distance of their difference from the 25^th^ or 75^th^ percentile exceeds 2.5 times the interquartile interval of their difference. Phase locking was tested for significance using the Rayleigh test for non-uniformity. Phase angles were compared using a circular non-parametric multi-sample test for equal medians. Differences in proportions were tested using Fisher’s exact test. Nested data were analyzed with linear mixed-effects models (LMEM) using animals as a fixed effect. Significance levels * p<0.05, ** p<0.01 and *** p<0.001 were considered. If not included in the text, values and corresponding test statistics of all presented data can be found in the supplementary material (Table S1-14).

## Acknowledgments

We thank A. Marquardt, A. Dahlmann, P. Putthoff and K. Titze for excellent technical assistance, Dr. I. Braren from the Vector Facility of the UKE for the virus production as well as Drs. M. Chini, S.H. Bitzenhofer, and R.L. van den Brink for helpful discussions. Moreover, we thank Dr. S.H. Bitzenhofer for building the olfactometer.

## Author Contributions

I.L.H.-O. and J.K.K conceived the study and designed the experiments. J.K.K carried out the experiments and analyzed the data. J.K.K, and I.L.H.-O. interpreted the data. J.K.K. and I.L.H.-O. wrote the article. All authors discussed and commented on the manuscript.

## Supplementary Figures

**S1 Fig. (related to Fig 1).**
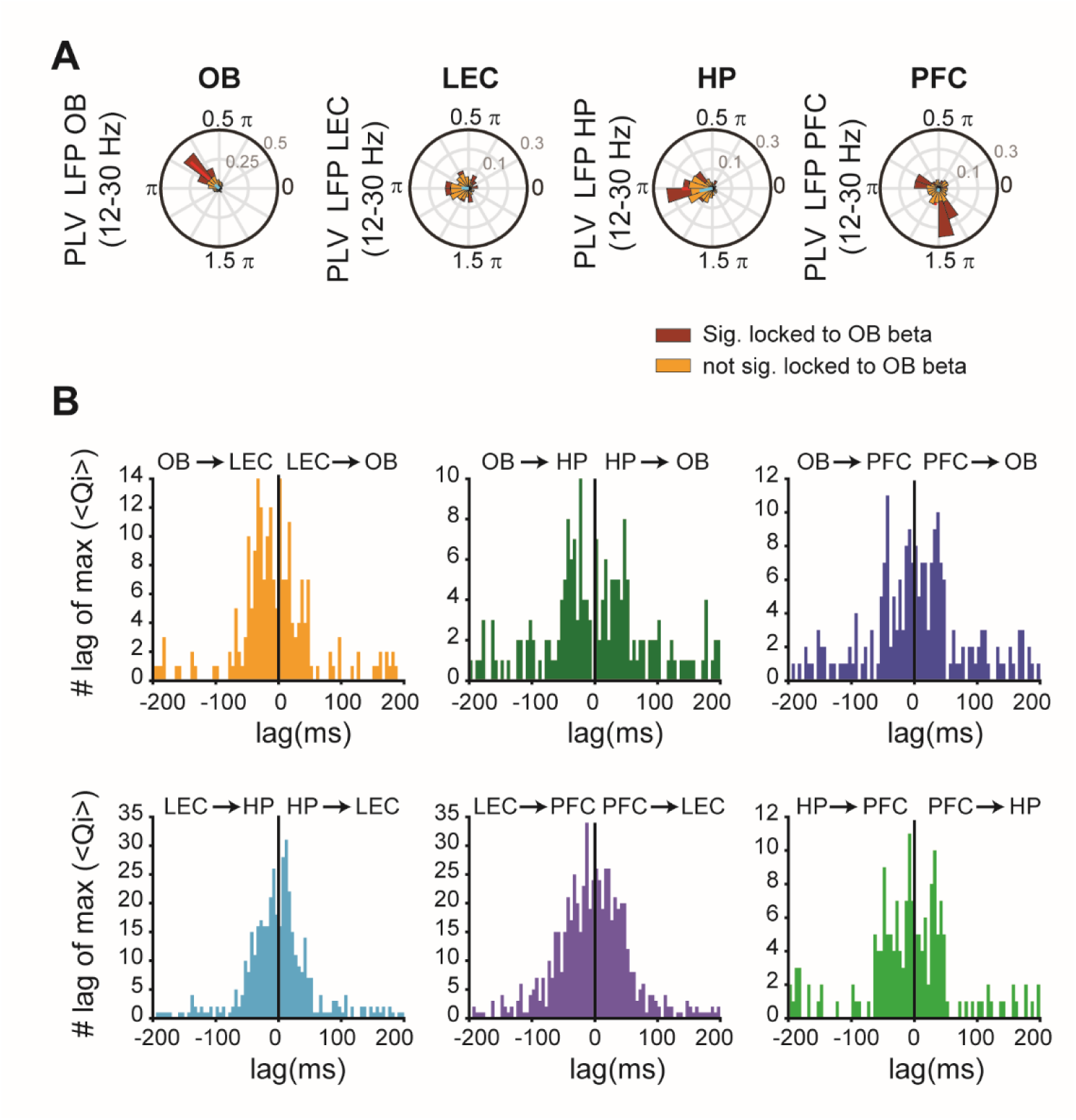
Phase locking of units to local beta oscillations and histograms of peak lag cross-covariance. **(A)** Polar plots displaying the phase-locking of units in OB, LEC, HP, and PFC to local beta oscillations for units that are (red) or are not (yellow) significantly locked to OB beta oscillations. **(B)** Histograms of peak lag of standardized mean spike-spike cross-covariance for OB-LEC (yellow), OB-HP (green), OB-PFC (blue), LEC-HP (light blue), LEC-PFC (purple), and HP-LEC (light green). Negative lags indicate that spiking in the first brain area precedes spiking in the second brain area.

**S2 Fig. (related to Fig 3).**
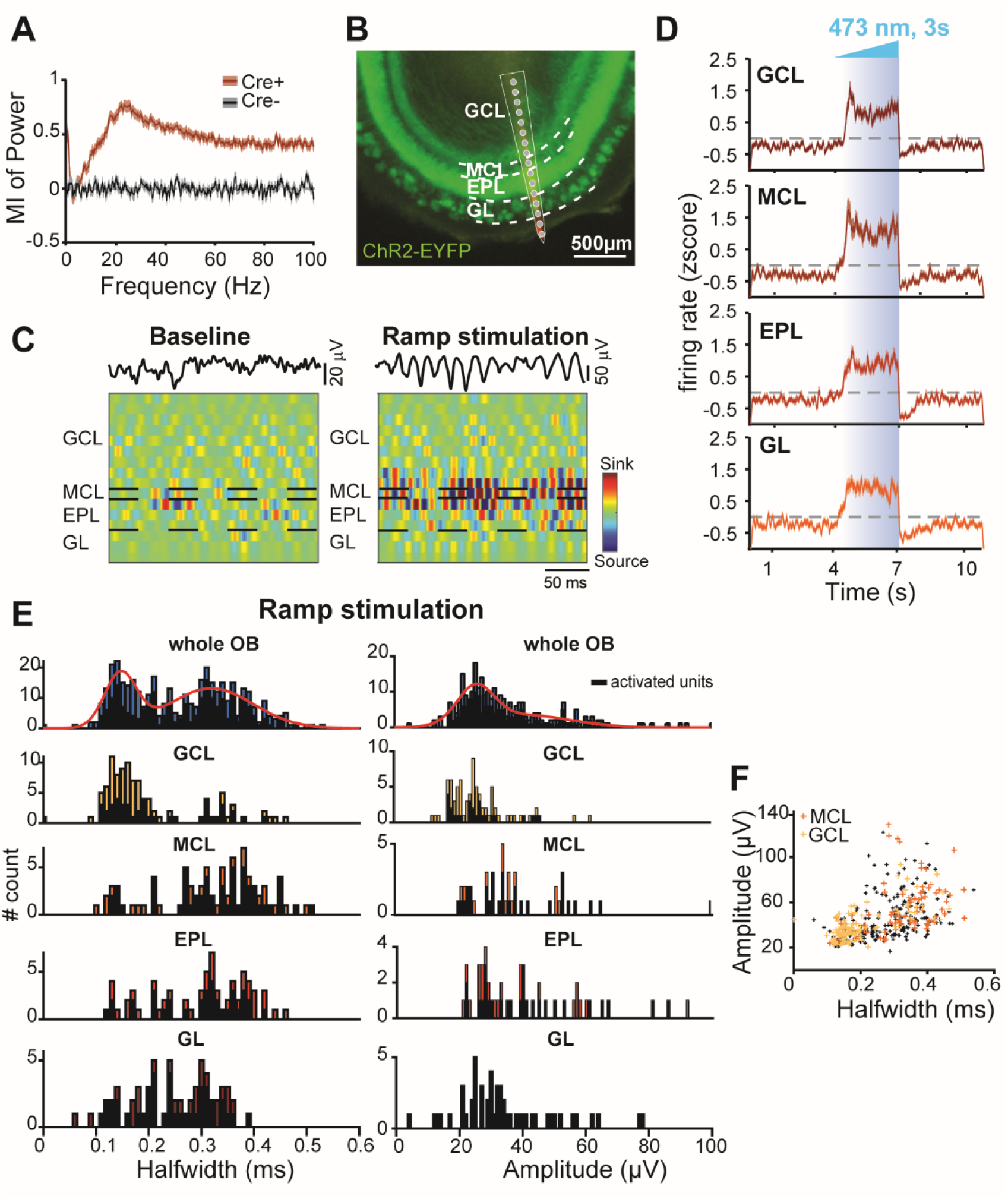
OB activity in response to ramp stimulation. **(A)** Plot of MI for LFP power in OB during ramp stimulation for cre^+^ (red) and cre^-^ (black) mice. **(B)** Digital photomontages displaying the DiI-labeled electrode track in a coronal slice including the ChR2-EYFP-expressing OB. Gray dots correspond to the individual recording sites. Dashed white lines mark borders between the different OB layers. **(C)** Filtered LFP traces (1-100 Hz) and current-source density profiles of spontaneous and light-evoked beta oscillations in the OB. **(D)** Z-scored firing rate of units recorded in the different OB layers in response to light stimulation. **(E)** Left: Histograms of waveform halfwidths of units recorded in the different layers of the OB. Right: same as left for amplitudes of unit waveforms. **(F)** Plot displaying the amplitudes and halfwidths of units recorded in the MCL (orange) and GCL (yellow) in the OB.

**S3 Fig. (related to Fig 3 and 4).**
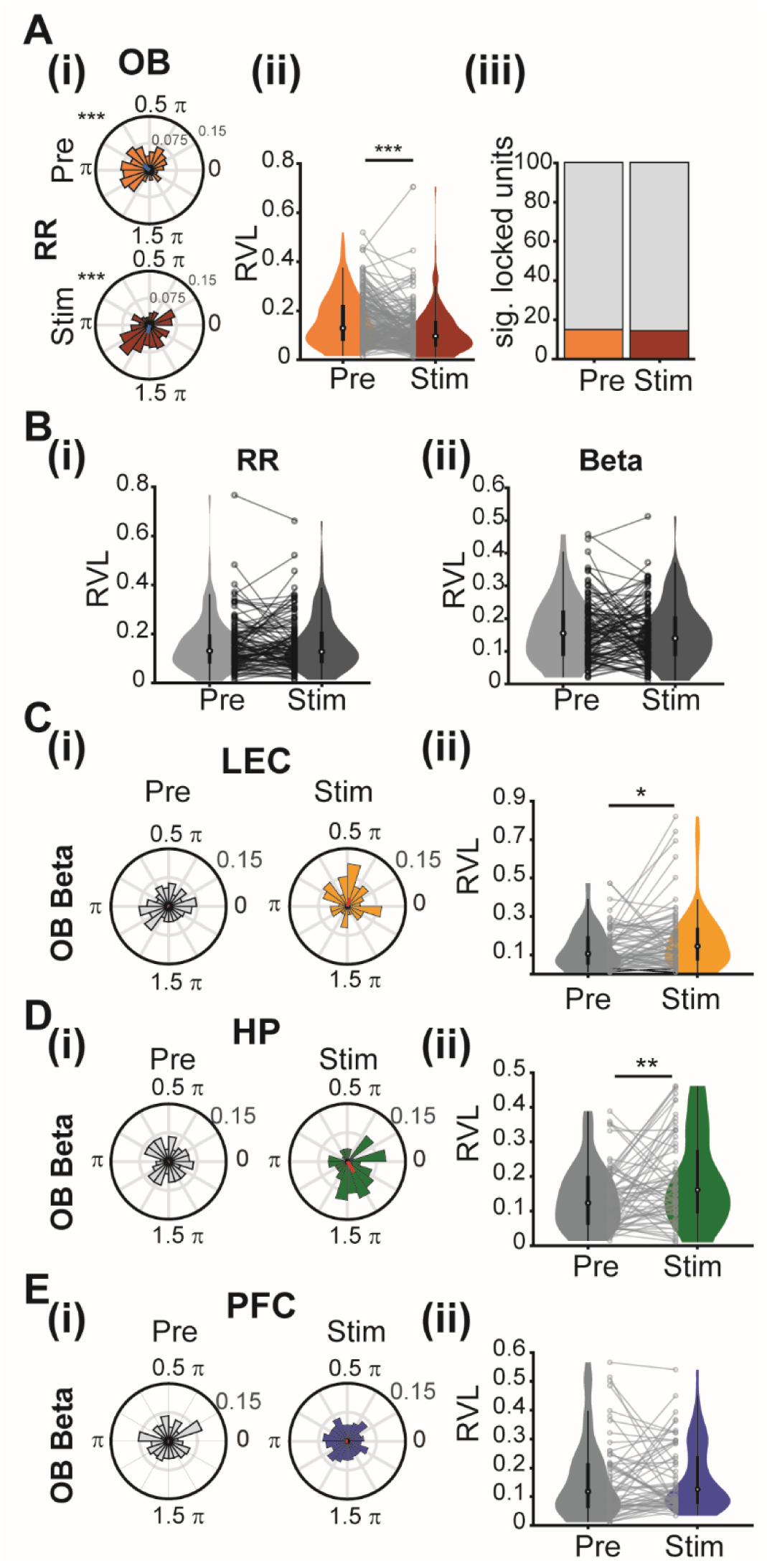
Phase locking of OB, LEC, HP, and PFC firing to beta oscillations in OB during the activation of MT/Cs by ramp stimulation. **(A) (i)** Polar plots displaying phase locking of OB units to RR before (Pre, orange) and during ramp stimulation (Stim, red) for cre^+^ mice. The mean resulting vectors are displayed as a blue line. (*** p < 0.001, Rayleigh test for non-uniformity). **(ii)** Violin plots displaying the RVL of OB units before (Pre, orange) and during ramp stimulation (Stim, red). Gray dots and lines correspond to individual units. (*** p < 0.001, LMEM). **(iii)** Bar plots displaying the percentage of significantly locked units before (Pre, orange) and during (Stim, red) stimulation. (* p < 0.05, Fisher’s exact test). **(B) (i)** Violin plots displaying the RVL of OB units locked to RR before stimulation (Pre, gray) and during ramp stimulation (Stim, black) for cre^-^ mice. Black dots and lines correspond to individual units. (* p < 0.05, linear mixed-effect model). **(ii)** Same as (i) for beta oscillations. **(C) (i)** Polar plots displaying phase locking of one example unit in LEC beta oscillations in OB before (Pre, gray) and during ramp stimulation (Stim, yellow) for cre^+^ mice. The mean resulting vectors are displayed as a red line. **(ii)** Violin plots displaying the RVL of LEC units before (Pre, gray) and during ramp stimulation (Stim, yellow). Gray dots and lines correspond to individual units. (* p < 0.05, ** p < 0.01, LMEM). **(D)** Same as (C) for HP. **(E)** Same as (C) for PFC.

**S4 Fig. (related to Figs 3 and 4).**
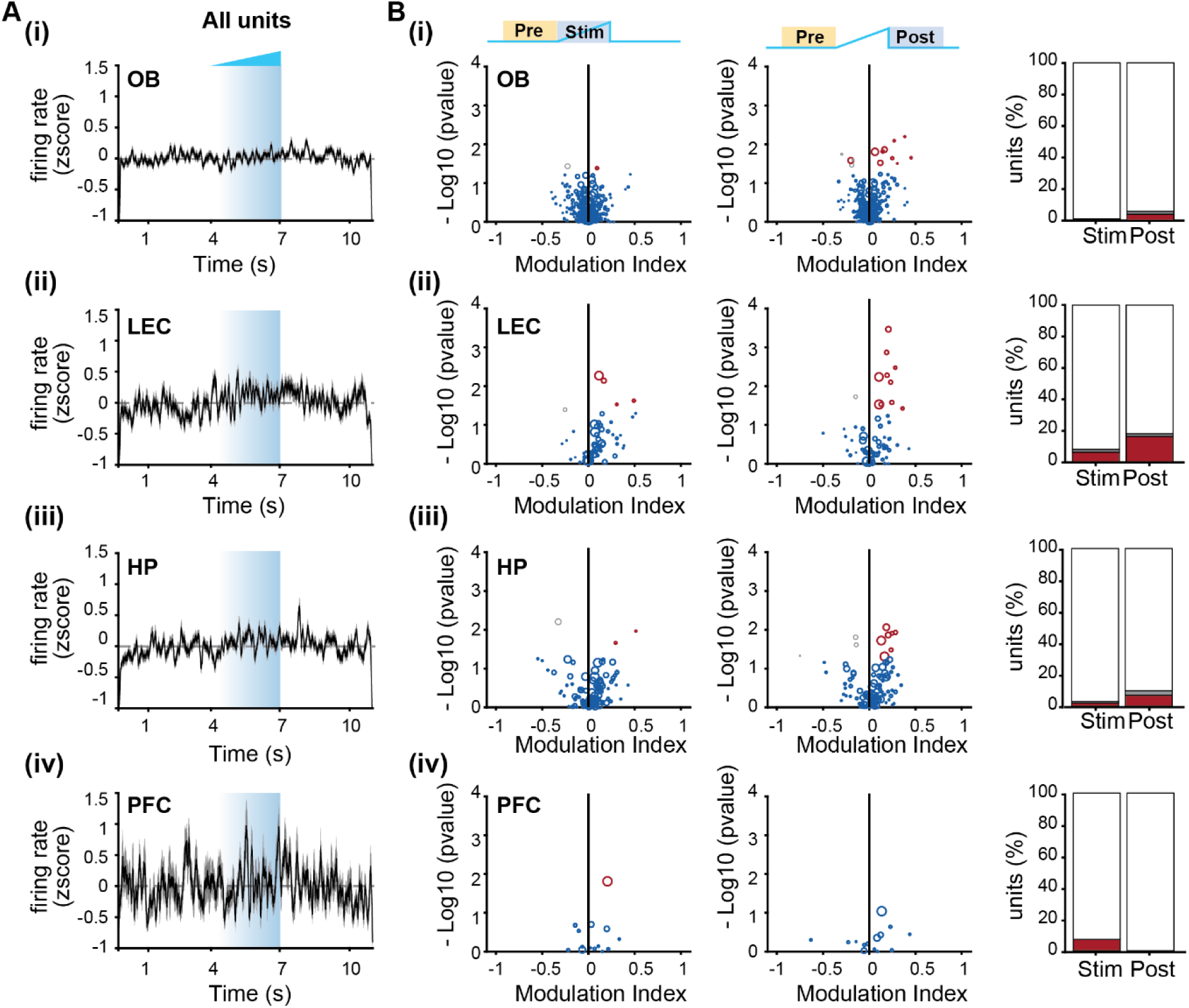
Effects of optogenetic manipulation of M/TCs from cre^-^ mice on single-unit activity. **(A) (i)** Z-scored firing rate of units recorded in the OB of cre^-^ (black) mice in response to light stimulation. **(ii)** Same as (i) for units recorded in LEC. **(iii)** Same as (i) for units recorded in HP. **(iv)** Same as (i) for units recorded in PFC. **(B)** (i) Left, volcano plot displaying the MI of SUA firing rates recorded in the OB of cre^-^ mice before (Pre) vs. during (Stim) ramp stimulation (significant activated units are shown in red and significant inhibited units in gray, p < 0.01, Wilcoxon signed-rank test). Middle, same as the left image but for SUA firing rates before (Pre) vs. after (Post) ramp stimulation. Right, bar plots depicting the percentage of activated (red) and inhibited (gray) units during (Stim) and after (Post) ramp stimulation. **(ii)** Same as (i) for units recorded in LEC. **(iii)** Same as (i) for units recorded in HP. **(iv)** Same as (i) for units recorded in PFC.

**S5 Fig. (related to Fig 6).**
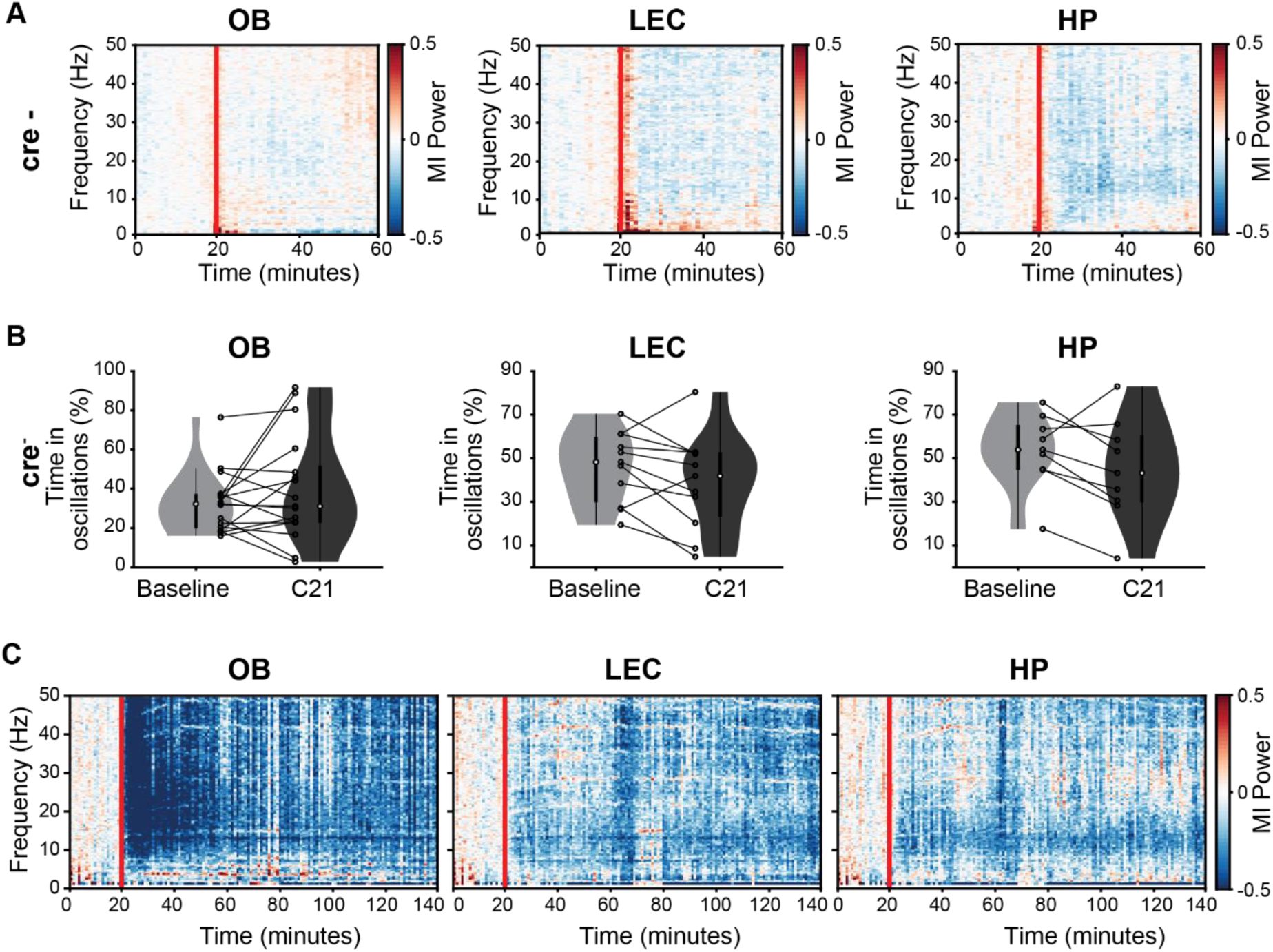
Effects of Compound 21 injection in cre^-^ mice and long-lasting effects of M/TC silencing by inhibitory DREADDs. **(A)** Color-coded MI of LFP power before and after C21 injection in OB (left), LEC (middle), and HP (right) of cre^-^ mice. The red line corresponds to the C21 injection. **(B)** Violin plots displaying the percentage of time spent in discontinuous oscillatory events before (Baseline, gray) and after C21 injection (C21, black) for cre^-^ mice. Black dots and lines correspond to individual animals. (* p < 0.05, Wilcoxon signed-rank test). **(C)** Color-coded MI of LFP power before and after (120 min) C21 injection in OB (left), LEC (middle) and HP (right) of cre^+^ mice (n=3). The red line corresponds to the C21 injection.

**S6 Fig. (related to Fig 6).**
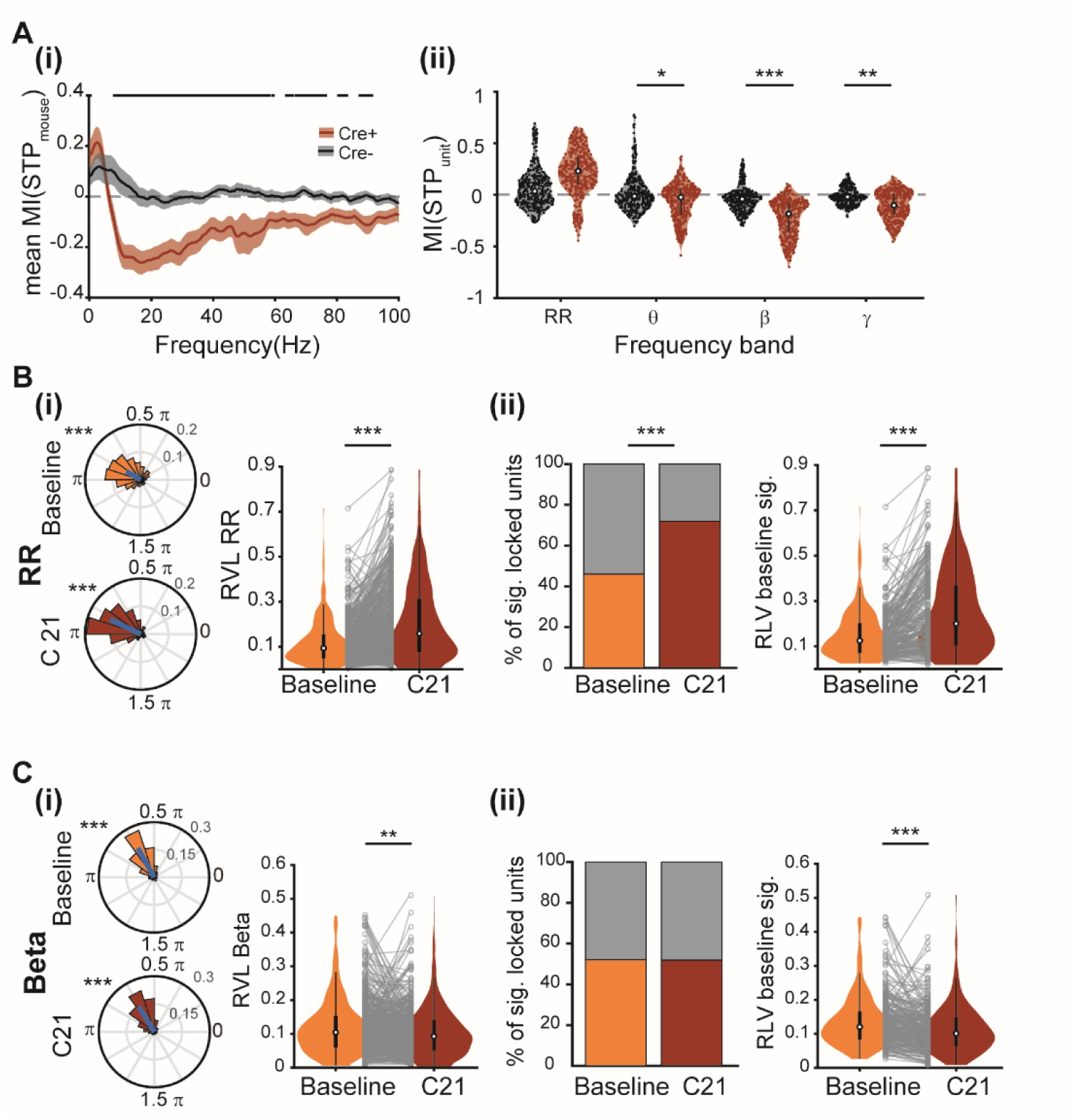
STP and phase locking of OB firing to LFP oscillations in OB after silencing the MT/C output by inhibitory DREADDs. **(A) (i)** Plot of MI of STP for cre^+^ (red) and cre^-^ (black) mice before and after C21 injection in OB. (black line: p < 0.05, Wilcoxon signed-rank test). **(ii)** Violin plots displaying mean MI for STP for each OB unit for different frequency bands for cre^+^ (red) and cre^-^ (black) mice. Red and black dots correspond to individual units. (* p < 0.05, ** p < 0.01, *** p < 0.001, linear mixed-effect model). **(B) (i)** Left, phase locking of OB units to RR oscillations in OB. Left, polar plots displaying phase locking of OB units before (baseline, orange) and after C21 injection (C21, red). The mean resulting vectors are represented as a blue line. (*** p < 0.001, Rayleigh test for non-uniformity). Right, violin plots displaying the RVL of OB units before (baseline, orange) and after C21 injection (C21, red). Gray dots and lines correspond to individual units. **(ii)** Left, bar plots displaying the percentage of significantly locked units before (baseline, yellow) and after (C21, red) C21 injection. (* p < 0.05, Fisher’s exact test). Right, violin plots displaying the RVL of OB units significantly locked to RR before C21 injection (baseline, orange) and the same units after C21 injection (C21, red). Gray dots and lines correspond to individual units. **(C)** Same as (B) for beta oscillations in the OB (** p < 0.01, *** p < 0.001, linear mixed-effect model).

**S7 Fig. (related to Fig 8).**
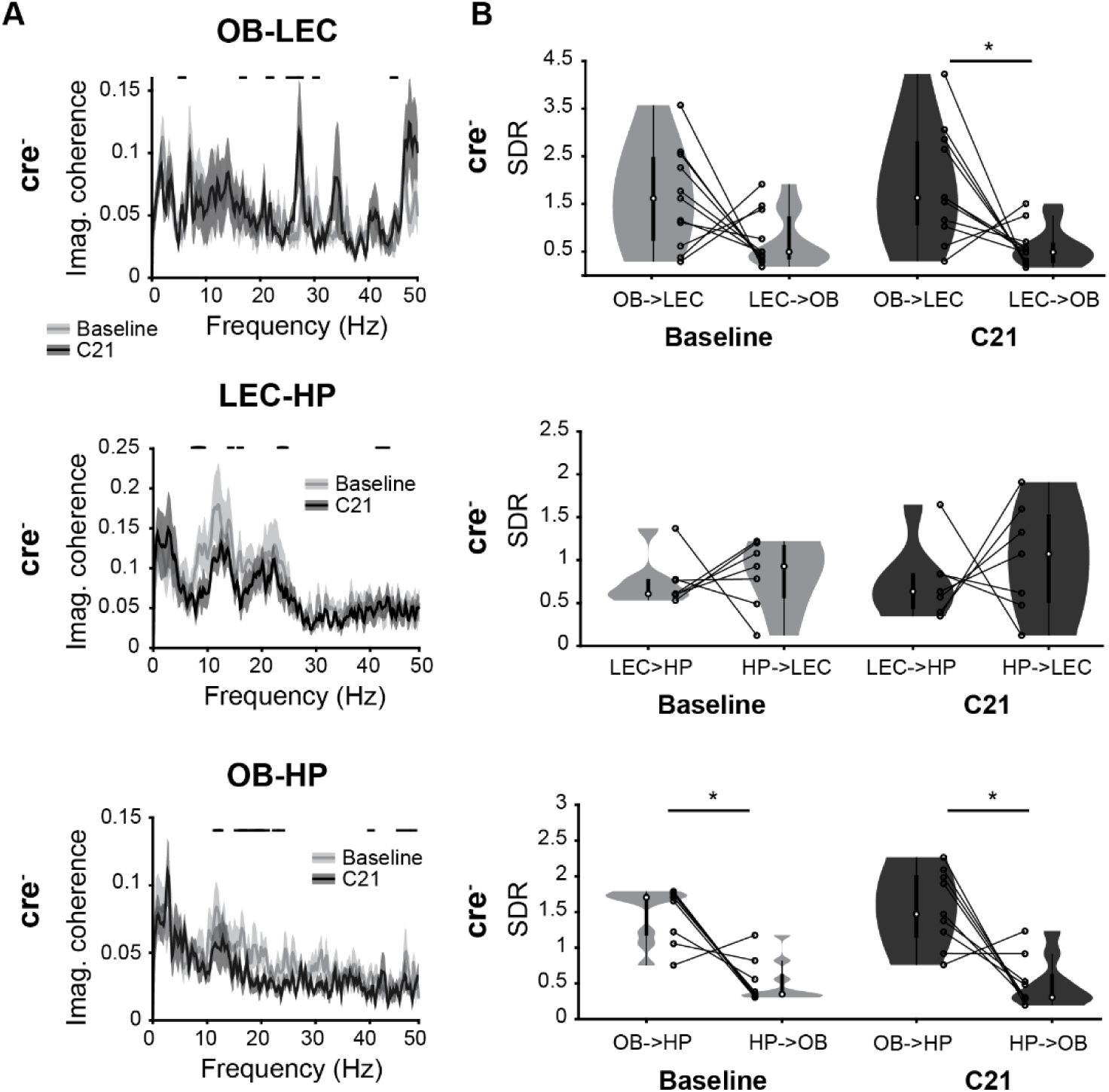
Effects of Compound 21 injection on oscillatory coupling in cre^-^ mice. **(A)** Imaginary coherence calculated for OB-LEC (top), LEC-HP (middle) and OB-HP (bottom) in cre^-^ mice, before (baseline, gray) and after C21 injection (C21, black). (black line: p < 0.05, Wilcoxon rank-sum test). **(B)** SDR for OB and LEC (top): SDR values for OB -> LEC and LEC -> OB before (baseline, gray) and after C21 injection (C21, black). Black dots and lines correspond to individual animals. Middle, same for LEC and HP. Bottom, same for OB and HP. (* p < 0.05, Wilcoxon signed-rank test).

## Supplementary Data Tables

**S1 Table (related to Fig 1D).** Phase locking of SUA to OB beta phase.

| | **Phase**<br>(circ. mean ± circ. SEM, % of sig.<br>locked units) | **p**<br>(Rayleigh test for non-uniformity) |
| --- | --- | --- |
| **OB** | 0.763 $\pi \pm 0.006 \pi$<br>640/1124 units (56.94 %) | $p=7.6*10^{-297}$ |
| **LEC** | 1.556 $\pi \pm 0.038 \pi$<br>42/384 units (10.94 %) | $p=4.19*10^{-11}$ |
| **HP** | 0.315 $\pi \pm 0.047 \pi$<br>51/384 units (13.28 %) | $p=3.68*10^{-5}$ |
| **PFC** | 1.715 $\pi \pm 0.082 \pi$<br>26/318 units (8.18 %) | $p=0.605$ |

**S2 Table (related to Fig 1D).**
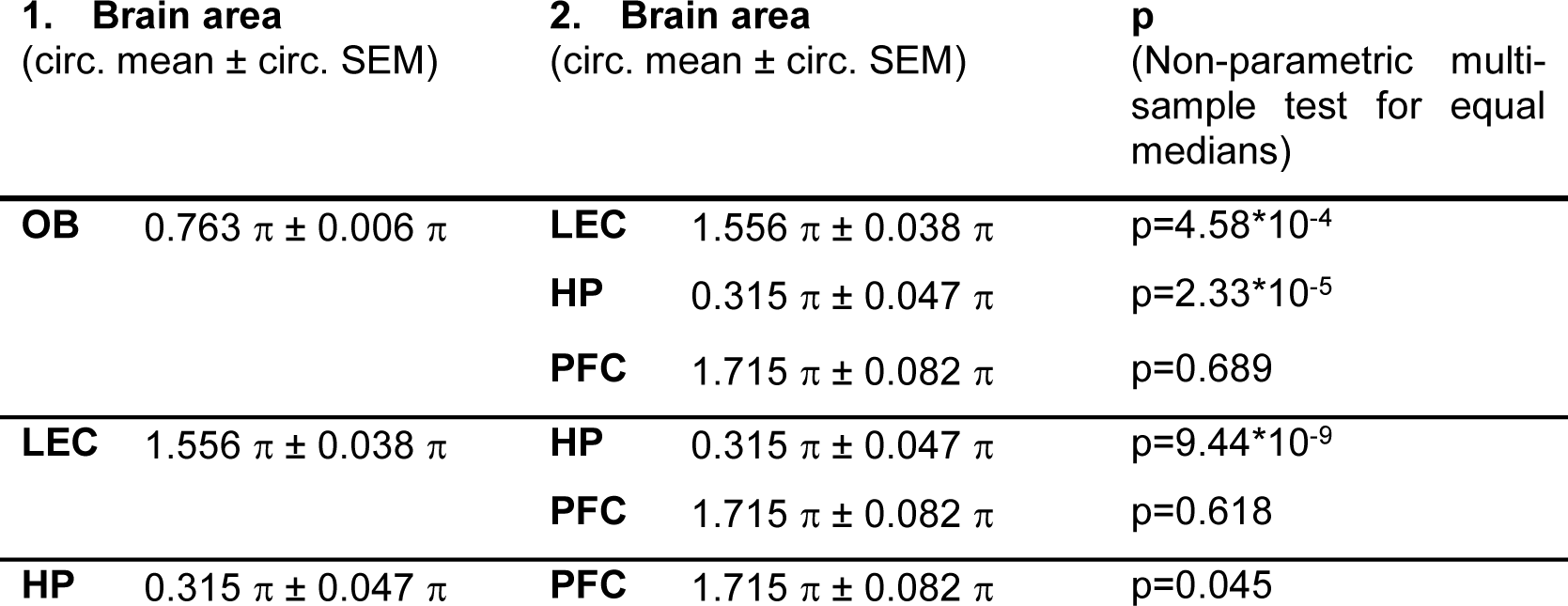
Phase angles of significant locked SUA to OB beta phase.

**S3 Table (related to Fig 1F).** SDR for pairs of brain areas.

|  | **Area 1 → Area 2**<br>(median, interquartile range) | **Area 2 → Area 1**<br>(median, interquartile range) | **p**<br>(Wilcoxon signed-rank test) |
| --- | --- | --- | --- |
| **Area 1: OB**<br>**Area 2: LEC** | 2.051 [1.666 – 2.580] | 0.407 [0.262 – 0.522] | p=5.68*10<sup>-8</sup> (n=39) |
| **Area 1: OB**<br>**Area 2: HP** | 1.389 [0.920 – 1.868] | 0.518 [0.334 – 0.657] | p=2.52*10<sup>-5</sup> (n=38) |
| **Area 1: OB**<br>**Area 2: PFC** | 2.809 [1.925 – 3.760] | 0.309 [0.226 – 0.399] | p=8.39*10<sup>-8</sup> (n=38) |
| **Area 1: LEC**<br>**Area 2: HP** | 0.650 [0.509 – 1.019] | 1.022 [0.831 – 1.372] | p=0.056 (n=28) |
| **Area 1: LEC**<br>**Area 2: PFC** | 1.188 [0.895 – 1.375] | 0.762 [0.635 – 1.045] | p=0.024 (n=30) |
| **Area 1: HP**<br>**Area 2: PFC** | 1.717 [1.317 – 2.034] | 0.481 [0.344 – 0.647] | p=0.0006 (n=24) |

**S4 Table (related to Fig 2E).** MI of power during olfactory stimulation.

|  |  | **MI of power**<br>(median, interquartile range) | **p**<br>Wilcoxon signed-rank test |
| --- | --- | --- | --- |
| **OB**<br>(n=18) | **RR (2-4 Hz)** | 0.451 [0.306 – 0.577] | p=1.96*10<sup>-4</sup> |
|  | **Theta (4-12 Hz)** | 0.481 [0.353 – 0.576] | p=1.96*10<sup>-4</sup> |
|  | **Beta (12-30 Hz)** | 0.472 [0.363 – 0.534] | p=1.96*10<sup>-4</sup> |
|  | **Gamma (30-100 Hz)** | 0.134 [0.089 – 0.187] | p=1.96*10<sup>-4</sup> |
| **LEC**<br>(n=11) | **RR (2-4 Hz)** | 0.185 [0.068 – 0.248] | p=0.007 |
|  | **Theta (4-12 Hz)** | 0.285 [0.229 – 0.327] | p=0.003 |
|  | **Beta (12-30 Hz)** | 0.258 [0.189 – 0.356] | p=0.002 |
|  | **Gamma (30-100 Hz)** | 0.089 [0.063 – 0.179] | p=9.77*10<sup>-4</sup> |
| **HP**<br>(n=18) | **RR (2-4 Hz)** | 0.060 [0.007 – 0.177] | p=0.022 |
|  | **Theta (4-12 Hz)** | 0.223 [0.107 – 0.294] | p=0.002 |
|  | **Beta (12-30 Hz)** | 0.251 [0.144 – 0.362] | p=0.002 |
|  | **Gamma (30-100 Hz)** | 0.081 [0.011 – 0.128] | p=0.035 |
| **PFC**<br>(n=14) | **RR (2-4 Hz)** | 0.256 [0.178 – 0.305] | p=2.44 *10<sup>-4</sup> |
|  | **Theta (4-12 Hz)** | 0.259 [0.203 – 0.308] | p=2.44 *10<sup>-4</sup> |
|  | **Beta (12-30 Hz)** | 0.252 [0.196 – 0.297] | p=2.44*10<sup>-4</sup> |
|  | **Gamma (30-100 Hz)** | 0.061 [0.045 – 0.090] | p=1.22 *10<sup>-4</sup> |

**S5 Table (related to Fig 2F).**
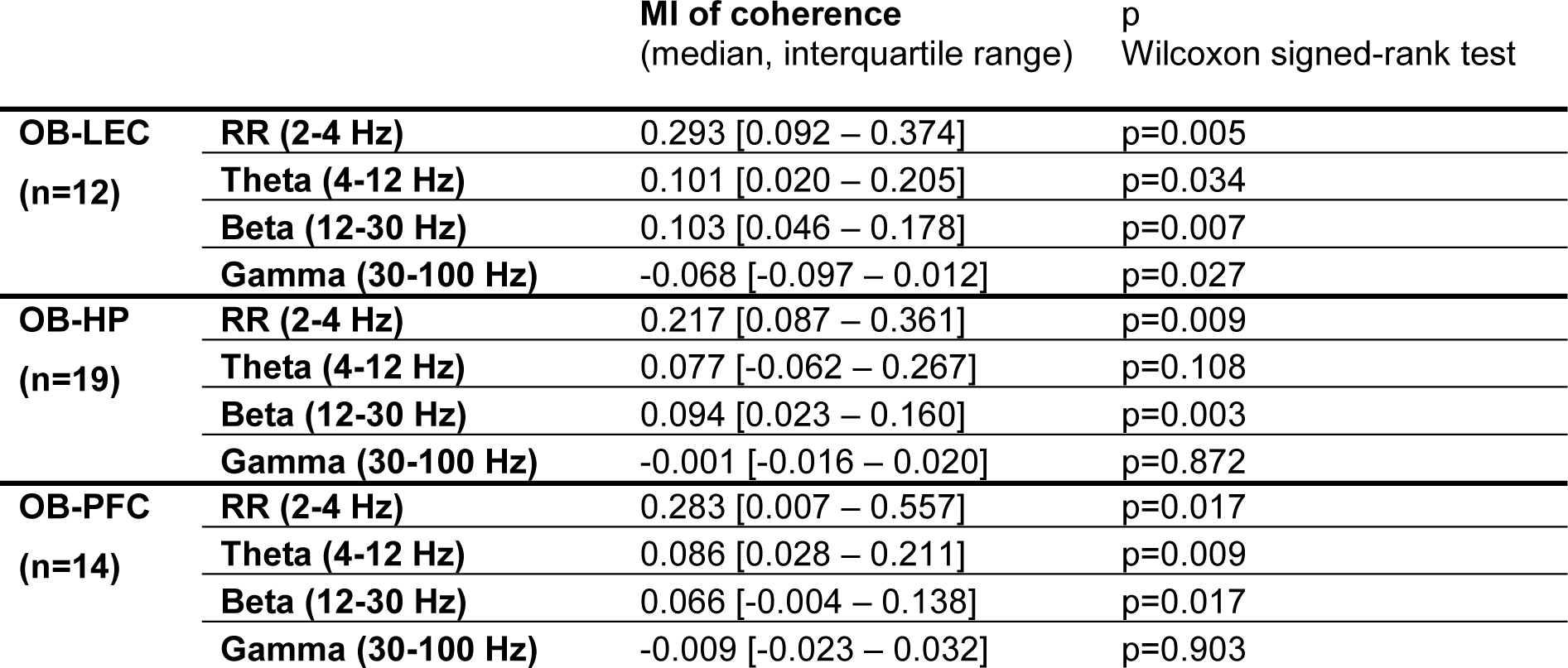
MI of coherence during olfactory stimulation.

|  |  | **MI of coherence**<br>(median, interquartile range) | p<br>Wilcoxon signed-rank test |
| --- | --- | --- | --- |
| **OB-LEC**<br>(n=12) | **RR (2-4 Hz)** | 0.293 [0.092 – 0.374] | p=0.005 |
|  | **Theta (4-12 Hz)** | 0.101 [0.020 – 0.205] | p=0.034 |
|  | **Beta (12-30 Hz)** | 0.103 [0.046 – 0.178] | p=0.007 |
|  | **Gamma (30-100 Hz)** | -0.068 [-0.097 – 0.012] | p=0.027 |
| **OB-HP**<br>(n=19) | **RR (2-4 Hz)** | 0.217 [0.087 – 0.361] | p=0.009 |
|  | **Theta (4-12 Hz)** | 0.077 [-0.062 – 0.267] | p=0.108 |
|  | **Beta (12-30 Hz)** | 0.094 [0.023 – 0.160] | p=0.003 |
|  | **Gamma (30-100 Hz)** | -0.001 [-0.016 – 0.020] | p=0.872 |
| **OB-PFC**<br>(n=14) | **RR (2-4 Hz)** | 0.283 [0.007 – 0.557] | p=0.017 |
|  | **Theta (4-12 Hz)** | 0.086 [0.028 – 0.211] | p=0.009 |
|  | **Beta (12-30 Hz)** | 0.066 [-0.004 – 0.138] | p=0.017 |
|  | **Gamma (30-100 Hz)** | -0.009 [-0.023 – 0.032] | p=0.903 |

**S6 Table (related to Figs 3D and 5B).** MI of power for cre^+^ and cre^-^ animals during light stimulation.

|  |  | **Cre<sup>+</sup>**<br>(median, interquartile<br>range, Wilcoxon signed-<br>rank test) | **Cre<sup>-</sup>**<br>(median, interquartile<br>range, Wilcoxon signed-<br>rank test) | **p**<br>(Wilcoxon<br>rank-sum test) |
| --- | --- | --- | --- | --- |
| **OB** | **RR (2-4 Hz)** | 0.045 [-0.047 – 0.139]<br>p=0.054 (n=32) | -0.010 [-0.067 – 0.045]<br>p=0.670 (n=14) | p=0.177 |
|  | **Theta (4-12 Hz)** | 0.184 [0.035 – 0.308]<br>p=5.81*10<sup>-5</sup> (n=32) | -0.014 [-0.025 – 0.031]<br>p=0.808 (n=14) | p=0.001 |
|  | **Beta (12-30 Hz)** | 0.746 [0.535 – 0.813]<br>p=7.95*10<sup>-7</sup> (n=32) | -0.002 [-0.014 – 0.028]<br>p=0.542 (n=14) | p=9.52*10<sup>-8</sup> |
|  | **Gamma (30-100 Hz)** | 0.555 [0.296 – 0.663]<br>p=7.95*10<sup>-7</sup> (n=32) | -0.007 [-0.038 – 0.021]<br>p=0.626 (n=14) | p=9.52*10<sup>-8</sup> |
| **LEC** | **RR (2-4 Hz)** | 0.093 [0.021 – 0.197]<br>p= 0.009 (n=22) | 0.083 [-0.040 – 0.153]<br>p=0.064 (n=10) | p=0.792 |
|  | **Theta (4-12 Hz)** | 0.089 [-0.011 – 0.182]<br>p= 0.005 (n=22) | 0.070 [0.017 – 0.130]<br>p=0.002 (n=10) | p=0.887 |
|  | **Beta (12-30 Hz)** | 0.107 [0.055 – 0.233]<br>p= 6.09*10<sup>-5</sup> (n=22) | 0.054 [-0.009 – 0.086]<br>p=0.084 (n=10) | p=0.033 |
|  | **Gamma (30-100 Hz)** | 0.0436 [0.016 – 0.104]<br>p=8.756*10<sup>-4</sup> (n=22) | 0.020 [-0.020 – 0.034]<br>p=0.432 (n=10) | p=0.064 |
| **HP** | **RR (2-4 Hz)** | -0.015 [-0.102 – 0.139]<br>p=0.858 (n=22) | 0.041 [-0.069 – 0.104]<br>p=0.4650 (n=11) | p=0.606 |
|  | **Theta (4-12 Hz)** | 0.052 [0.019 – 0.167]<br>p=0.001 (n=22) | 0.023 [-0.033 – 0.101]<br>p=0.520 (n=11) | p=0.122 |
|  | **Beta (12-30 Hz)** | 0.135 [0.032 – 0.202]<br>p=4.83*10<sup>-4</sup> (n=22) | 0.026 [-0.046 – 0.079]<br>p=0.520 (n=11) | p=0.021 |
|  | **Gamma (30-100 Hz)** | 0.068 [0.016 – 0.114]<br>p= 0.001 (n=22) | 0.040 [-0.026 – 0.059]<br>p=0.206 (n=11) | p=0.089 |
| **PFC** | **RR (2-4 Hz)** | 0.065 [-0.011 – 0.177]<br>p=0.016 (n=22) | 0.016 [-0.038 – 0.119]<br>p=0.461 (n=8) | p=0.439 |
|  | **Theta (4-12 Hz)** | 0.081 [-0.008 – 0.155]<br>p=0.01 (n=22) | 0.035 [-0.01 – 0.062]<br>p=0.25 (n=8) | p=0.386 |
|  | **Beta (12-30 Hz)** | 0.120 [0.093 – 0.327]<br>p=4.613*10<sup>-5</sup> (n=22) | 0.005 [-0.050 – 0.025]<br>p=0.641 (n=8) | p=2.78*10<sup>-4</sup> |
|  | **Gamma (30-100 Hz)** | 0.061 [-0.007 – 0.082]<br>p=0.002 (n=22) | 0.012 [-0.013 – 0.063]<br>p=0.383 (n=8) | p=0.291 |

**S7 Table (related to S3B Fig).** Mean resulting vector length for phase locking of OB firing to RR and beta oscillations for cre^-^ mice.

| **Cre-**<br>(n<sub>units</sub>=100<br>13 mice) | **Pre**<br>(median, interquartile<br>range) | **Stim**<br>(median, interquartile<br>range) | **p**<br>(linear fixed-<br>effect model) |
| --- | --- | --- | --- |
| **RR (2-4 Hz)** | 0.131 [0.083 – 0.196] | 0.128 [0.085 – 0.208] | p=0.763 |
| **Beta (12-30 Hz)** | 0.155 [0.088 – 0.222] | 0.140 [0.087 – 0.205] | p=0.486 |

**S8 Table (related to Fig 5C).** MI of coherence for cre^+^ and cre^-^ animals during light stimulation.

|  |  | **Cre<sup>+</sup>**<br>(median, interquartile<br>range, Wilcoxon<br>signed-rank test) | **Cre<sup>-</sup>**<br>(median, interquartile<br>range, Wilcoxon signed-<br>rank test) | **p**<br>(Wilcoxon<br>rank-sum<br>test) |
| --- | --- | --- | --- | --- |
| **OB-LEC** | **RR (2-4 Hz)** | 0.053 [-0.166 – 0.184]<br>p=0.59 (n=21) | -0.031 [-0.128 – 0.197]<br>p=0.922 (n=10) | p=0.751 |
|  | **Theta (4-12 Hz)** | -0.013 [-0.134 – 0.119]<br>p=0.543 (n=21) | 0.038 [-0.024 – 0.108]<br>p=0.375 (n=10) | p=0.386 |
|  | **Beta (12-30 Hz)** | 0.129 [-0.009 – 0.182]<br>p=0.005 (n=21) | 0.014 [-0.016 – 0.035]<br>p=0.695 (n=10) | p=0.036 |
|  | **Gamma (30-100 Hz)** | 0.01 [-0.062 – 0.039]<br>p=0.639 (n=21) | 0.003 [-0.019 – 0.037]<br>p=0.922 (n=10) | p=0.949 |
| **OB-HP** | **RR (2-4 Hz)** | 0.001 [-0.106 – 0.070]<br>p=0.741 (n=21) | 0.059 [-0.092 – 0.136]<br>p=0.638 (n=11) | p=0.5 |
|  | **Theta (4-12 Hz)** | 0.010 [-0.028 – 0.114]<br>p= 0.394 (n=21) | -0.034 [-0.151 – 0.042]<br>0.365 (n=11) | p=0.177 |
|  | **Beta (12-30 Hz)** | 0.074 [0.021 – 0.134]<br>p=0.005 (n=21) | -0.053 [-0.123 – 0.063]<br>p=0.365 (n=11) | p=0.017 |
|  | **Gamma (30-100 Hz)** | -0.020 [-0.049 – 0.015]<br>p=0.149 (n=21) | 0.025 [-0.026 – 0.039]<br>p=0.52 (n=11) | p=0.177 |
| **OB-PFC** | **RR (2-4 Hz)** | -0.022 [-0.090 – 0.135]<br>p=0.689 (n=21) | -0.113 [-0.208 – 0.123]<br>p=0.641 (n=8) | p=0.542 |
|  | **Theta (4-12 Hz)** | -0.053 [-0.098 – 0.076]<br>p=0.543 (n=21) | -0.035 [-0.178 – 0.120]<br>p=0.844 (n=8) | p=0.942 |
|  | **Beta (12-30 Hz)** | 0.159 [0.026 – 0.265]<br>p=0.0004 (n=21) | -0.010 [-0.123 – 0.085]<br>p=0.742 (n=8) | p=0.008 |
|  | **Gamma (30-100 Hz)** | 0.037 [0.009 – 0.088]<br>p=0.01 (n=21) | -0.025 [-0.055 – 0.029]<br>p=0.641 (n=8) | p=0.067 |

**S9 Table (related to Fig 6F).** MI of power after C21 injection.

|  |  | **Cre<sup>+</sup>**<br>(median, interquartile<br>range, Wilcoxon signed-<br>rank test) | **Cre<sup>-</sup>**<br>(median, interquartile<br>range, Wilcoxon signed-<br>rank test) | **p**<br>(Wilcoxon<br>rank-sum test) |
| --- | --- | --- | --- | --- |
| **OB** | **RR (2-4 Hz)** | -0.032 [-0.160 – 0.157]<br>p=0.687 (n=17) | -0.025 [-0.084 – 0.116]<br>p=0.879 (n=18) | p=0.609 |
|  | **Theta (4-12 Hz)** | -0.082 [-0.286 – -0.014]<br>p=0.013 (n=17) | -0.010 [-0.094 – 0.156]<br>p=0.472 (n=18) | p=0.031 |
|  | **Beta (12-30 Hz)** | -0.246 [-0.426 – -0.117]<br>p=0.0003 (n=17) | -0.025 [-0.137 – 0.052]<br>p=0.349 (n=18) | p=0.0003 |
|  | **Gamma (30-100 Hz)** | -0.162 [-0.225 – 0.006]<br>p=0.006 (n=17) | -0.009 [-0.083 – 0.040]<br>p=0.647 (n=18) | p=0.028 |
| **LEC** | **RR (2-4 Hz)** | -0.337 [-0.492 – -0.109]<br>p=0.002 (n=13) | 0.246 [-0.096 – 0.488]<br>p=0.063 (n=12) | p=0.001 |
|  | **Theta (4-12 Hz)** | -0.293 [-0.435 – -0.10]<br>p= 0.001 (n=13) | 0.080 [-0.126 – 0.330]<br>0.259 (n=12) | p=0.0008 |
|  | **Beta (12-30 Hz)** | -0.195 [-0.424 – -0.115]<br>p=0.0002 (n=13) | -0.043 [-0.106 – 0.025]<br>p=0.226 (n=12) | p=0.0005 |
|  | **Gamma (30-100 Hz)** | -0.106 [-0.151 – -0.044]<br>p=0.0005 (n=13) | -0.015 [-0.050 – 0.323]<br>p=0.663 (n=12) | p=0.004 |
| **HP** | **RR (2-4 Hz)** | -0.131 [-0.285 – -0.047]<br>p=0.049 (n=10) | -0.074 [-0.160 – 0.036]<br>p=0.426 (n=9) | p=0.243 |
|  | **Theta (4-12 Hz)** | -0.202 [-0.332 – -0.144]<br>p=0.006 (n=10) | -0.053 [-0.116 – 0.017]<br>p=0.359 (n=9) | p=0.010 |
|  | **Beta (12-30 Hz)** | -0.221 [-0.288 – -0.150]<br>p=0.004 (n=10) | -0.069 [-0.112 – 0.001]<br>p=0.055 (n=9) | p=0.017 |
|  | **Gamma (30-100 Hz)** | -0.072 [-0.119 – -0.013]<br>p=0.027 (n=10) | 0.003 [-0.043 – 0.033]<br>p=1 (n=9) | p=0.065 |

**S10 Table (related to Fig 6G and S5B Fig).** Time in discontinuous events.

|  |  | **Baseline**<br>(median, interquartile<br>range, in %) | **Compound 21**<br>(median, interquartile<br>range, in %) | **p**<br>(Wilcoxon signed-rank test) |
| --- | --- | --- | --- | --- |
| **OB** | **Cre<sup>+</sup> (n=17)** | 26.131 [20.490 – 32.841] | 19.083 [2.638 – 24.127] | p=0.031 |
|  | **Cre<sup>-</sup> (n=17)** | 32.310 [20.032 – 37.206] | 31.103 [22.888 – 51.465] | p=0.435 |
| **LEC** | **Cre<sup>+</sup> (n=13)** | 48.216 [35.999 – 53.788] | 14.889 [6.576 – 28.939] | p=0.0005 |
|  | **Cre<sup>-</sup> (n=11)** | 48.294 [29.988 – 59.585] | 41.896 [23.372 – 52.576] | p=0.147 |
| **HP** | **Cre<sup>+</sup> (n=10)** | 40.596 [36.197 – 53.448] | 18.349 [9.143 – 33.924] | p=0.002 |
|  | **Cre<sup>-</sup> (n=9)** | 53.911 [44.874 – 64.950] | 43.233 [30.020 – 60.254] | p=0.164 |

**S11 Table (related to S6A Fig).** STP for OB after C21 injection.

|  |  | **Cre<sup>+</sup>**<br>(median, interquartile<br>range) | **Cre<sup>-</sup>**<br>(median, interquartile<br>range) | **p**<br>(linear mixed-<br>effect model) |
| --- | --- | --- | --- | --- |
|  |  | n<sub>units</sub>=415 from 15 mice | n<sub>units</sub>=361 from 16 mice |  |
| **OB** | **RR (2-4 Hz)** | 0.230 [0.096 – 0.367] | 0.036 [-0.074 – 0.160] | p=0.351 |
|  | **Theta (4-12 Hz)** | -0.025 [-0.189 – 0.056] | -0.014 [-0.098 – 0.093] | p=0.017 |
|  | **Beta (12-30 Hz)** | -0.184 [-0.368 – -0.075] | -0.041 [-0.121 – 0.009] | p=0.0002 |
|  | **Gamma (30-100 Hz)** | -0.103 [-0.195 – 0.016] | -0.020 [-0.071 – 0.044] | p=0.005 |

**S12 Table (related to Fig 8B).** MI coherence for cre^+^ and cre^-^ mice after C21 injection.

|  |  | **Cre<sup>+</sup>**<br>(median, interquartile<br>range, Wilcoxon signed-<br>rank test) | **Cre<sup>-</sup>**<br>(median, interquartile<br>range, Wilcoxon signed-<br>rank test) | **p**<br>(Wilcoxon<br>rank-sum<br>test) |
| --- | --- | --- | --- | --- |
| **OB-LEC** | **RR (2-4 Hz)** | -0.089 [-0.217 – 0.156]<br>p= 0.774 (n=12) | -0.103 [-0.368 – 0.212]<br>p=1 (n=11) | p=0.976 |
|  | **Theta (4-12 Hz)** | -0.085 [-0.473 – 0.226]<br>p=1 (n=12) | -0.074 [-0.218 – 0.083]<br>p=0.549 (n=11) | p=0.976 |
|  | **Beta (12-30 Hz)** | -0.165 [-0.340 – 0.050]<br>p= 0.039 (n=12) | 0.005 [-0.119 – 0.226]<br>p=0.549 (n=11) | p=0.021 |
|  | **Gamma (30-100 Hz)** | -0.018 [-0.104 – 0.162]<br>p= 0.774 (n=12) | -0.021 [-0.124 – 0.330]<br>p =1 (n=11) | p=0.735 |
| **LEC-HP** | **RR (2-4 Hz)** | -0.072 [-0.268 – 0.528]<br>p=0.727 (n=8) | 0.024 [-0.089 – 0.251]<br>p=1 (n=7) | p=0.867 |
|  | **Theta (4-12 Hz)** | -0.082 [-0.436 – 0.092]<br>p= 0.727 (n=8) | -0.176 [-0.304 – -0.056]<br>0.125 (n=7) | p=0.955 |
|  | **Beta (12-30 Hz)** | -0.095 [-0.214 – -0.026]<br>p=0.070 (n=8) | -0.147 [-0.211 – -0.024]<br>p=0.125 (n=7) | p=0.779 |
|  | **Gamma (30-100 Hz)** | -0.079 [-0.183 – 0.133]<br>p=0.727 (n=8) | -0.023 [-0.053 – 0.090]<br>p=1 (n=7) | p=0.536 |
| **OB-HP** | **RR (2-4 Hz)** | 0.311 [0.214 – 0.501]<br>p=0.109 (n=10) | 0.169 [-0.303 – 0.212]<br>p=1 (n=9) | p=0.182 |
|  | **Theta (4-12 Hz)** | 0.045 [-0.174 – 0.341]<br>p=0.754 (n=10) | -0.047 [-0.204 – -0.006]<br>p=0.180 (n=9) | p=0.356 |
|  | **Beta (12-30 Hz)** | -0.318 [-0.397 – -0.124]<br>p=0.021 (n=10) | -0.178 [-0.306 – -0.058]<br>p=0.039 (n=9) | p=0.278 |
|  | **Gamma (30-100 Hz)** | 0.036 [-0.067 – 0.161]<br>p=1 (n=10) | -0.074 [-0.096 – 0.060]<br>p=0.508 (n=9) | p=0.095 |

**S13 Table (related to Fig 8D and S7B Fig).**
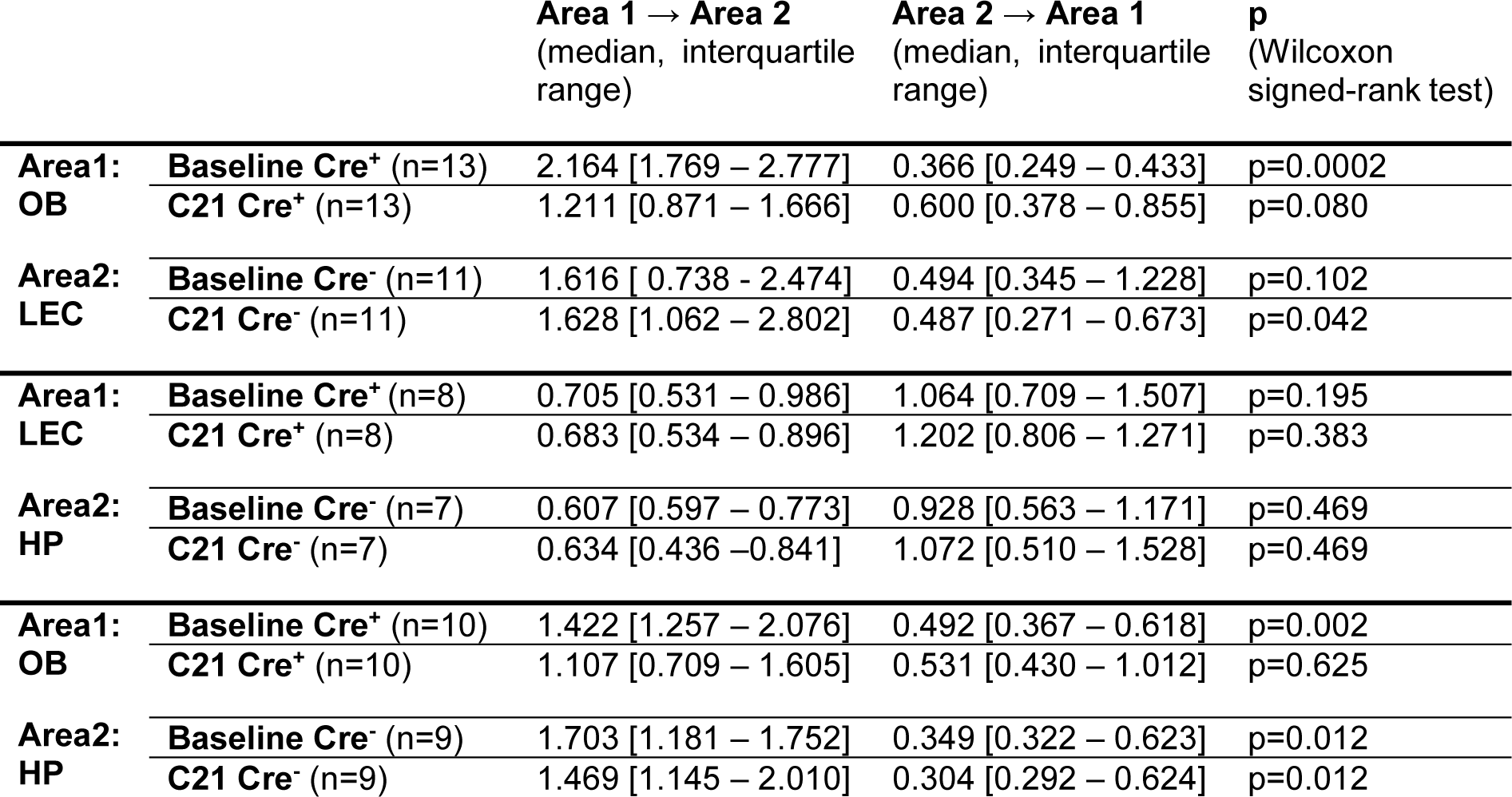
SDR before and after C21 injection.

**S14 Table (related to Fig 8D and S7B Fig).** SDR difference for before and after C21 injection.

|  |  | **Baseline**<br>(median, interquartile range) | **Compound 21**<br>(median, interquartile range) | **p**<br>(Wilcoxon signed-rank test) |
| --- | --- | --- | --- | --- |
| **Area1: OB**<br>**Area2: LEC** | **Difference Cre<sup>+</sup> (n=13)** | 1.792 [1.300 – 2.512] | 0.619 [0.087 – 1.160] | p=0.001 |
|  | **Difference Cre<sup>-</sup> (n=11)** | 1.192 [-0.490 – 2.130] | 1.045 [0.479 – 2.531] | p=0.175 |
| **Area1: LEC**<br>**Area2: HP** | **Difference Cre<sup>+</sup> (n=8)** | -0.357 [-0.929 – 0.254] | -0.471 [-0.746 – 0.041] | p=0.641 |
|  | **Difference Cre<sup>-</sup> (n=7)** | -0.326 [-0.582 – 0.210] | -0.438 [-1.092 – 0.331] | p=0.813 |
| **Area1: OB**<br>**Area2: HP** | **Difference Cre<sup>+</sup> (n=10)** | 0.891 [0.601 – 1.709] | 0.627 [-0.303 – 1.175] | p=0.004 |
|  | **Difference Cre<sup>-</sup> (n=9)** | 1.347 [0.557 – 1.413] | 1.167 [0.553 – 1.708] | p=0.496 |

